# A modular patient-derived organoid–xenograft platform reveals molecular and clinical trajectories of prostate cancer progression

**DOI:** 10.64898/2026.01.05.697006

**Authors:** Romuald Parmentier, Robin Dolgos, Raphaëlle Servant, Jing Wang, Luca Roma, Renaud Mevel, Daniela Bossi, Heike Pueschel, Zoi Diamantopoulou, Tatjana Vlajnic, Frank Stenner, Arnoud J. Templeton, Helge Seifert, Nicola Aceto, Jean-Phillipe Theurillat, Cyrill A. Rentsch, Lukas Bubendorf, Clémentine Le Magnen

**Affiliations:** Pathology, Institute of Medical Genetics and Pathology, University Hospital Basel, University of Basel, Switzerland; Department of Urology, University Hospital Basel, Basel, Switzerland; Department of Biomedicine, University of Basel, University Hospital Basel, Basel, Switzerland; Institute of Oncology Research, Bellinzona & Faculty of Biomedical Sciences, Università della Svizzera italiana (USI), Lugano, Switzerland; Department of Biology, Institute of Molecular Health Sciences, Swiss Federal Institute of Technology (ETH) Zurich, Switzerland; Division of Medical Oncology, University Hospital Basel, Basel, Switzerland; Division of Medical Oncology, St. Claraspital, Basel, and Faculty of Medicine, University of Basel, Basel, Switzerland

## Abstract

Patient-derived organoids (PDOs) are becoming increasingly important in prostate cancer (PCa) translational research. However, direct proof-of-concept studies demonstrating their ability to model disease evolution, identify relevant biomarkers, and accurately predict treatment response in PCa patients, remain scarce. Here, we report the establishment of serially transplantable xenografts series derived from two advanced PCa PDO lines, which can be further re-cultured as organoids. Newly-generated model series maintain key phenotypic, genomic, and functional characteristics of the original patient tumors, and emulate relevant molecular subtypes of advanced PCa. Single-cell RNA sequencing (scRNA-seq) analysis uncovers transcriptomic differences between xenograft and organoid models, as well as signaling pathways which are largely preserved and can be targeted *ex vivo.* Functional drug profiles correlate with molecular and clinical attributes, as exemplified by response to androgen receptor (AR) pathway inhibitors and glucocorticoid-mediated AR signaling activation. Longitudinal scRNA-seq analysis of perturbed PDOs identifies a rare PROX1^+^/ALDH1A1^+^ cell population, which pre-exist in the treatment-naïve setting and is significantly enriched upon androgen deprivation. Notably, this cell population is similarly enriched in post-treatment samples of the original patient and is associated with aggressive AR-negative PCa molecular subtypes, suggesting a potential link with PCa progression. Our study provides proof-of-concept evidence that organoids can mirror PCa patient-specific drug sensitivity profiles and molecular paths of disease progression, uncovering pertinent biomarkers. Ultimately, our organoid-xenograft model series provide a modular and scalable platform that can readily be used for mechanistic and translational studies.

## Introduction

Prostate cancer (PCa) is among the top most diagnosed cancer type and leading cause of cancer-related deaths in European men (*1, 2*). Improving survival of men with advanced disease remains an important unmet clinical need (*3*). In these men, androgen deprivation therapy (ADT i.e., surgical or chemical castration) has been the mainstay of treatment for several decades. Although most patients initially respond well to ADT and the disease is referred to as hormone-sensitive (HSPC), the majority will eventually relapse and progress to a castration-resistant state (CRPC) which currently remains incurable. While the treatment landscape for advanced PCa is rapidly evolving (*4, 5*), predictive biomarkers that could guide therapy in these patients are emerging but are still scarce (*6–8*). In addition, mechanisms underlying the transition from advanced HSPC to CRPC, and ultimately to aggressive CRPC subtypes such as neuroendocrine PCa (NEPC) and those negative for both androgen receptor (AR) and neuroendocrine (NE) markers (i.e., double-negative PCa; DNPC) (*9*), are still poorly understood. To address these clinical challenges, there is still an urgent need for preclinical models that (i) emulate the advanced hormone-sensitive state and have the capacity to progress to more aggressive disease states, and (ii) derive from patients resistant to recently approved therapies and recapitulate clinical response/resistance profiles (*10*).

Progress in this field has long been hindered by a lack of experimental model systems that capture the molecular and cellular complexity of PCa. Illustrating this issue, PCa patient-derived xenografts (PDXs) still only represent 0.17% of all xenografts in international collections despite the high prevalence of PCa worldwide (*10, 11*). This low number mainly reflects the poor engraftment rates associated with the establishment of PCa PDXs and a limited access to fresh samples with aggressive phenotypes. More recently, patient-derived organoids (PDOs) have emerged as an alternative to PDXs; despite recent advancements, the success rates associated with the generation of clinically relevant expandable PCa PDO models remain, however, unsatisfactory (*12–17*). Furthermore, although PDOs are increasingly used to test drug responses, direct proof-of-concept studies demonstrating their ability to model disease evolution and uncover insights into treatment resistance relevant to the clinical setting are lacking.

In this study, we report the establishment of two serially transplantable xenografts series derived from PCa patient-derived organoid cultures (referred to as PDOXs) which can be further re-cultured as organoids (referred to as PDOXOs). Newly-generated model series maintain the histological, genomic and functional characteristics of the original patient tumors, and model relevant molecular subtypes of advanced PCa. Through single-cell RNA sequencing (scRNA-seq) comparison of PDOX and PDOXO models, we reveal transcriptomic differences and signaling pathways which are largely preserved *in vitro* and can be targeted in functional drug testing experiments. Leveraging longitudinal scRNA-seq analyses, we identify minor tumor cell subpopulations that are enriched upon androgen-deprivation in PDOs and in the post-treatment patient’ samples. We provide proof-of-concept evidence that organoids can be used not only to test drug responses but also to model clinical trajectories observed in the matched patients, thereby uncovering patient-specific paths of disease progression.

## Results

### Generation of serially transplantable patient-derived organoid xenograft models of advanced prostate cancer

Two patient-derived organoid cultures previously generated in our laboratory were used to generate PDOX models (*14*). P20-11 PDOs were derived from a lung metastasis obtained from a patient with treatment-naïve metastatic HSPC (mHSPC) (**Figure 1a**, and methods). P20-23 PDOs were derived from a transurethral resection of the prostate (TURP) obtained from a patient with metastatic CRPC (mCRPC). The latter had previously been treated with a combination of ADT (goserelin) and docetaxel, and was under treatment with the AR pathway inhibitor (ARPi) enzalutamide for 11 months, at the time of resection (**Figure 1a**, and methods).

**Figure 1.**
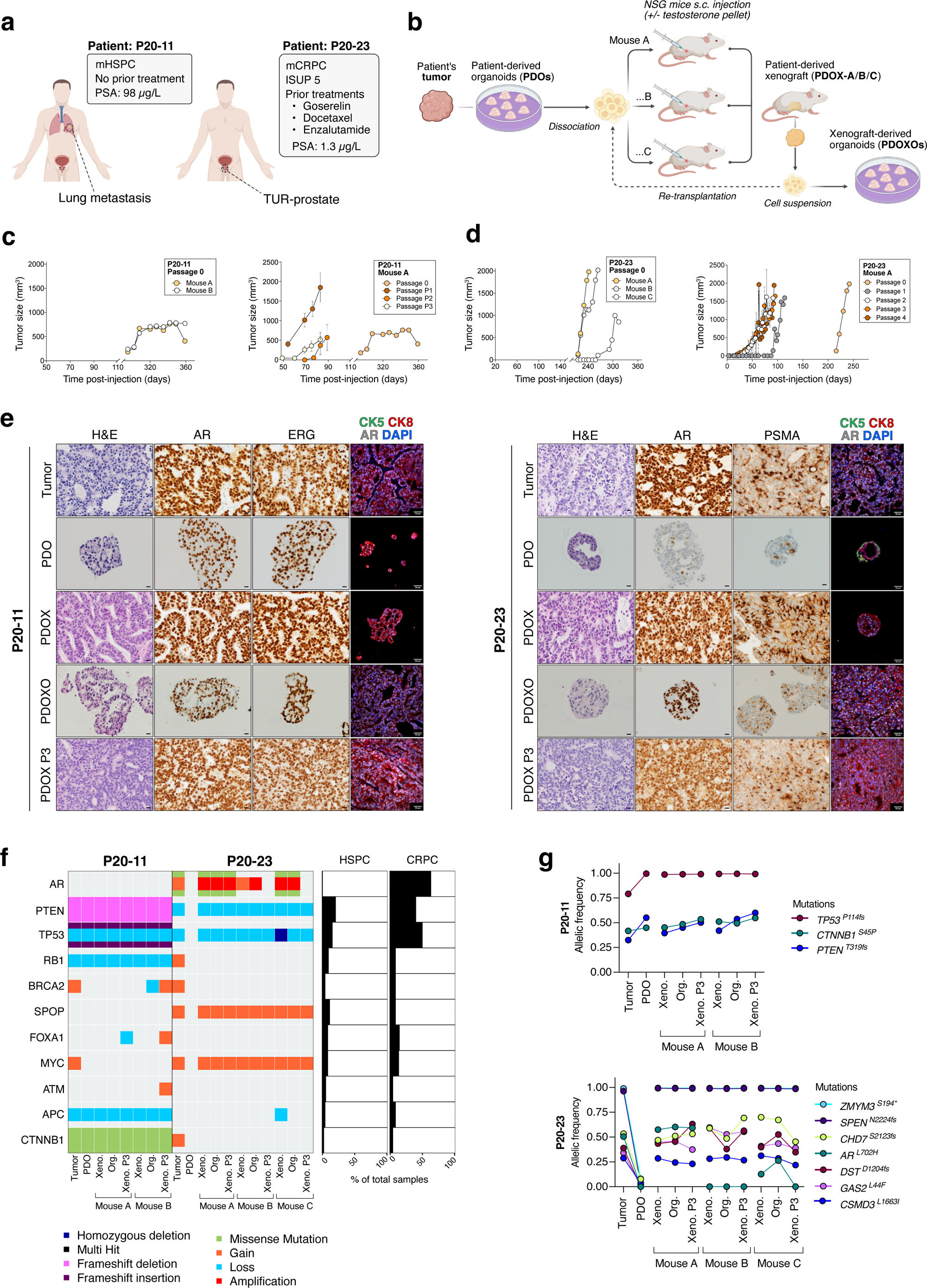
Generation of two novel serially-transplantable patient-derived model series representative of advanced prostate cancer. **(a)** Cartoons depicting the clinical background of patients from whom P20-11 and P20-23 samples were obtained. mHSPC: metastatic Hormone Sensitive Prostate Cancer, PSA: Prostate Specific Antigen, mCRPC: metastatic Castration Resistant Prostate Cancer. **(b)** Experimental workflow used to derive patient-derived organoids (PDOs), PDO-derived xenografts (PDOXs) and PDOX-derived organoids (PDOXOs). s.c.: subcutaneous; NSG: NOD scid gamma. **(c)** P20-11 PDOX tumor growth kinetics at initial implantation (PDOX-A & B, left) and later passages (PDOX-A, right). Tumor growth kinetics of PDOX-B over serial passaging is represented in Figure S1. **(d)** P20-23 PDOX tumor growth kinetics at initial implantation (PDOX-A, B & C, left) and later passages (PDOX-A, right). PDOX tumor growth kinetics of PDOX-B & C over serial passaging are represented in Figure S1. **(e)** Phenotypic analyses of patient tumors and matched PDOs, PDOX, PDOXOs, and PDOX P3 for P20-11 (PDOX-B, left panel) and P20-23 (PDOX-C, right panel). Shown are representative H&E images, immunofluorescence (IF) and Immunohistochemistry (IHC) staining for the indicated antibodies. Scale bars represent 20 µM for IHC/H&E images and 50 µM for IF images. **(f) Left**: Oncoplot depicting selected PCa-relevant genomic alterations in P20-11 and P20-23 model series as assessed by whole exome sequencing. **Right**: genomic alteration frequency in a metastatic HSPC cohort (*20*) and in a CRPC cohort (*21*) represented as % of total samples. Xeno: Xenograft, Org: Organoid. **(g)** Allelic frequency analysis of relevant non-synonymous mutations detected in P20-11 and P20-23 model series.

To ensure the long-term preservation and expansion of both PDO lines, organoid-derived cells were injected into the flank of male NOD *scid* gamma mice with or without testosterone pellets (P20-11 and P20-23, respectively; **Figure 1b; Table S1**). Engraftment rates and growth kinetics varied between both models and passages (**Figure 1c-d; Figure S1a**). At each xenograft generation, tumors were dissociated and used to derive organoids referred to as PDOXOs (**Figure 1b**). Interestingly, over generations *in vivo*, P20-11 PDOXOs grew for extended time and passages, suggesting that they may have acquired changes promoting their growth *in vitro* (**Figure S1b)**. In contrast, the first P20-23 PDOX generation had limited growth as PDOXOs and usually stopped at passage 2-3.

### PDOX and PDOXO models emulate relevant molecular subtypes of advanced PCa

*In vivo* and *in vitro* environments have been suggested to induce selection and/or the emergence of specific phenotypic and genomic clones (*18, 19*). We therefore characterized the features of our models using H&E, immunohistochemistry (IHC), immunofluorescence (IF), and whole exome sequencing (WES). Importantly, all PDOXs and PDOXOs were confirmed to be of human origin, as indicated by their positivity for human mitochondria (**Figure S1c**).

P20-11 patient tumor and PDOs displayed a purely luminal CK8_+_ phenotype with strong positivity for the androgen receptor (AR) and ERG, as previously reported (*14*) (**Figure 1e, left**). These features were maintained through *in vivo* generations and *in vitro* passages (passage 0 and 3). A high degree of similarity in the copy number variation (CNV) and mutation profiles was observed between all models (**Figure S1d-f**). In particular, mutations in genes commonly altered in mHSPC patients (*20*), such as *CTNNB1, PTEN,* and *TP53,* were detected with similar allelic frequencies (AF); furthermore, heterozygous deletions in *TP53* and *RB1,* as well as loss of PTEN expression were found in all the samples (**Figure 1f-g**; **Figure S1h)**. P20-23 original tumor displayed strong AR and PSMA expression, as well as a CK8+ luminal phenotype (**Figure 1e, right**). In contrast, P20-23 PDOs were characterized by a majority of cells with a benign phenotype (AR− PSMA− CK5+) and a minor subset of cells with a luminal cancer phenotype (AR+ PSMA+ CK5-), from which the xenografts may have derived. In line with this hypothesis, PDOX and PDOXO samples displayed a phenotype mirroring that of the original tumor (i.e., strong AR, PSMA, and CK8 expression, loss of PTEN; **Figure S1h**). Consistently, P20-23 PDOs carried mutations with low AF and shared only 37% of mutations with the original tumor while the PDOX-A xenograft shared 83% of mutations with the patient sample (**Figure S1f**). Mutations were mostly maintained at similar AF between the tumors and the PDOX models regardless of the PDOX subline, with the exception of the missense *AR_L702H_* mutation (AF: ∼50% in PDOX-A and tumor vs. ∼25% in PDOX-C vs. absent in PDOX-B; **Figure 1f-g, Figure S1g-h**). In addition to this pathogenic mutation, an *AR* amplification was also detected at different extents depending on the subline and in *vitro/in vivo* setting. Notably, AR alterations represent the most frequent genomic alterations in CRPC patients ((*21*) and **Figure 1f**). Taken together, these findings highlight that P20-11 and P20-23 model series preserve key molecular features of their original tumors and are representative of advanced HSPC and CRPC clinical cohorts, respectively.

### Transcriptomic heterogeneity and fidelity of models at single cell resolution

PDOXs and PDOXOs are frequently used as complementary models in translational cancer research (*22*); however, the degree of transcriptomic fidelity and heterogeneity across passages, as well as the relevance of putative transcriptomic differences between such model types, have not been investigated at single-cell resolution yet. To address these questions, we performed single-cell RNA sequencing (scRNA-seq) at distinct passages of P20-11/P20-23 PDOX and PDOXO models, when technically feasible. The UMAP projection clustered P20-11 and P20-23 model series separately on the UMAP1 axis, while the UMAP2 axis distinguished between *in vivo* and i*n vitro* counterparts, independently from cell cycle states (**Figure 2a, Figure S2a**). For each patient-derived model series, PDOX tumors and PDOXOs clustered together respectively, suggesting limited transcriptomic changes associated with *in vivo* or *in vitro* passaging.

**Figure 2.**
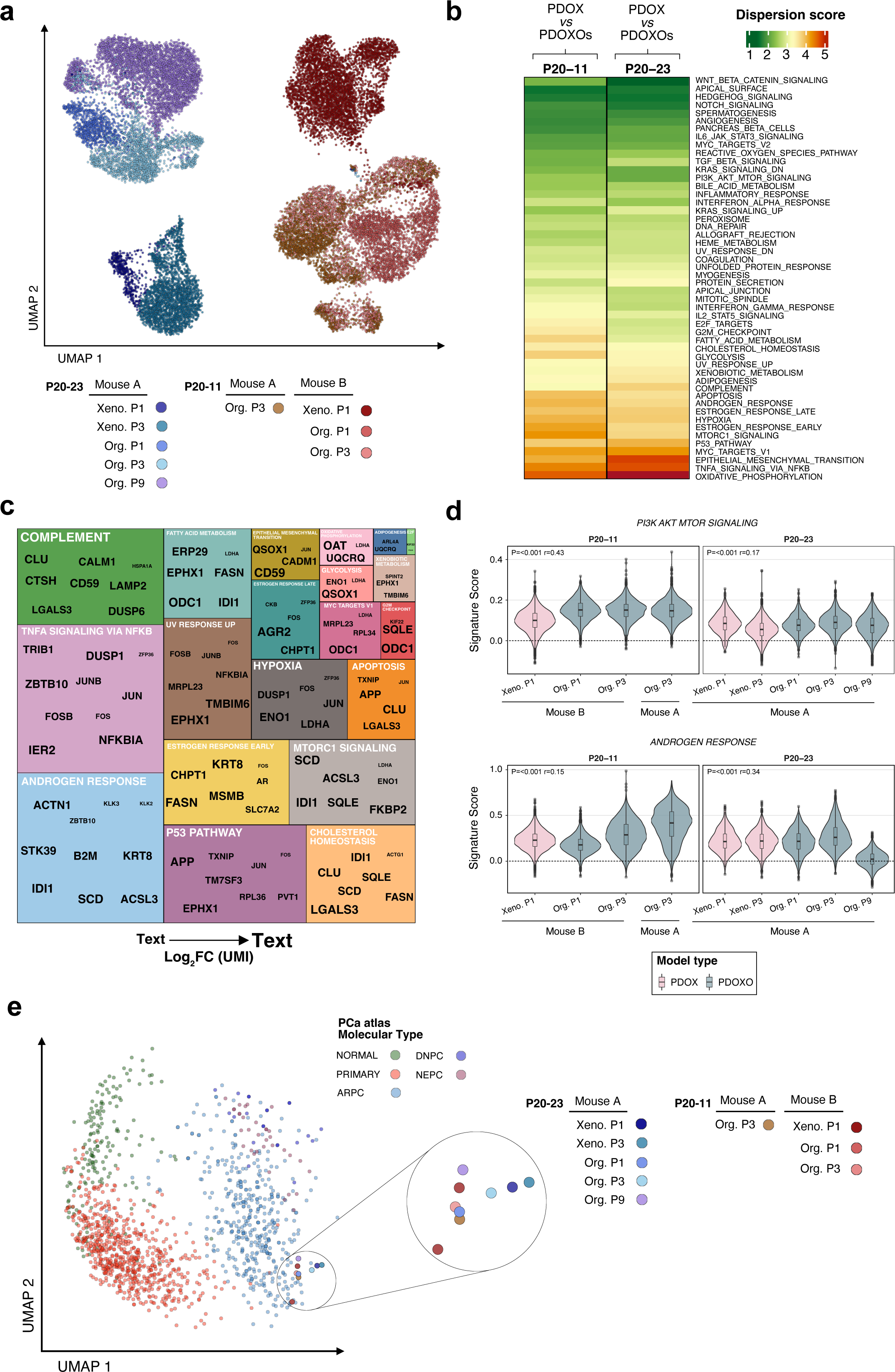
Single-cell RNA sequencing analysis of the models and *in vitro* transcriptomic shifts. **(a)** UMAP projection of PDOX and PDOXO cells color-coded by series, model and passages. **(b)** Per–series visualization of pathway dispersion scores computed between models (PDOXs vs PDOXOs). Dispersion scores were calculated using Pagoda2 on MSigDB Hallmark gene sets and pathways were ranked according to their average dispersion score across both series. **(c)** Treemap representation of the top 20 most dispersed pathways (according to average). Rectangle areas represent number of significantly differentially expressed genes above logFC 1.5 threshold within the selected pathways. Font size of individual genes represents values of log2 fold change relative to maximum fold change within each pathway. **(d)** Violin plots representing signature scores distribution across series, colored by model type (PDOXs or PDOXOs). Top violin plots represent *PI3K-AKT-MTOR SIGNALING* signature (MsigDB, Hallmark). Bottom violin plots represent *ANDROGEN RESPONSE* (MsigDB, Hallmark). Statistical analysis was performed using the Wilcoxon signed-rank test to compare PDOX and PDOXO groups. All reported P values were significant (P < 0.001). Effect sizes (r) were interpreted as follows: r < 0.2 (small), r < 0.5 (medium), r < 0.8 (substantial). Note that the small effect sizes, despite statistically significant differences, indicate broadly comparable activity of the signatures across models (i.e., PDOXs vs PDOXOs). **(e)** UMAP projection of the prostate cancer atlas (*23*), where each dot represents a patient, colored according to molecular subtype. ARPC: androgen receptor–positive PCa; DNPC: double-negative PCa; NEPC: neuroendocrine PCa. Integrated pseudo-bulk scRNA-seq profiles from our model series are highlighted.

To identify pathways putatively altered between *in vivo* (xenograft) and *in vitro* (organoid) counterparts, we next performed pathway overdispersion analysis (Pagoda2) among the HALLMARKS signature set (MSigDB) within each model series. Notably, both the list of pathways and their magnitude were comparable between PDOXs and PDOXOs in P20-11 and P20-23 model series (**Figure 2b**). Among the most dispersed pathways were those linked to central regulators such as P53 or MYC, to nuclear hormone receptors such as the Androgen and the Estrogen responses, and to metabolic states (i.e., OXIDATIVE PHOSPHORYLATION, HYPOXIA, GLYCOLYSIS). Of note, the three most dispersed pathways were “Epithelial to Mesenchymal transition”, “TNF alpha signaling via NFKB” and “Oxidative phosphorylation”.

To identify genes driving the partial pathway dispersion observed between PDOXs and PDOXOs, we performed differential expression analysis between these two settings, independently from patient origin, and narrowed down the analysis to the genes in the top 20 most dispersed pathways. When selecting the genes with the highest fold-change, the most affected pathways were “TNFA via NFKB” and “Androgen Response” (**Figure 2c**). Within the ANDROGEN RESPONSE signature, we found major downstream elements such as KLK2 and KLK3, which we previously identified as downregulated in PCa PDOs as compared to patient samples (*12*). In the “TNFA VIA NFKB” signature, genes with a central role in cell proliferation, apoptosis and inflammation such as JUNB, JUN, FOSB and FOS, were represented. While this analysis highlighted that pathway-specific transcriptional rewiring may occur at the gene level between the *in vitro* and *in vivo* contexts, the small effect sizes observed for the corresponding pathway signature scores indicate that core signaling programs, including “PI3K/AKT/MTOR” and “Androgen Response,” remain largely preserved (**Figure 2d**). This suggests that despite localized transcriptional shifts, PDOX and PDOXO models preserve the functional integrity of major oncogenic programs, making them suitable for mechanistic and therapeutic studies targeting these pathways.

To place our newly-generated models amongst a transcriptional landscape of PCa, we mapped their transcriptomic profiles with PCa clinical samples and existing experimental models using the PCa profiler tool (*23*) (**Figure 2e**). Consistent with an active AR signaling axis, all our models clustered within the AR+ PCa (ARPC) category, close to the commonly used androgen-sensitive LuCaP and LNCaP PDX model series, as well as the more recent PNPCa xenograft model (*15, 24*) (**Figure S2b)**.

### P20-11 and P20-23 model series are androgen-dependent *in vitro* and *in vivo*

Given their strong AR expression and sustained AR signaling activity, we next investigated whether our models were sensitive to androgen deprivation. PDOXOs were cultured with DHT (1 nM, androgen-proficient) or without DHT (0 nM, androgen-deprived) and cell viability was measured after 1, 2, and 3 weeks of culture in the two conditions (**Figure 3a**). After 7 days, the viability of P20-11 PDOXOs was significantly decreased in androgen-deprived conditions (one-way ANOVA, p ≤ 0.01), while no significant differences were observed in P20-23 PDOXOs. The most striking difference between the models was observed after 21 days in culture, whereby the relative viability was 5% for P20-11 and 40% for P20-23 (one-way ANOVA, p ≤ 0.0001) and accompanied by clear organoid size reduction in androgen-deprived conditions (**Figure 3b**). In line with these data, P20-11 and P20-23 PDOXs responded to surgical castration *in vivo* with high efficacy (7/7 tumors for P20-11; 5/5 tumors for P20-23; **Figure 3c**). Notably, no spontaneous regrowth was observed in castrated conditions for any of the models (up to 200 days). However, PDOXs were capable of regrowth following testosterone implantation, albeit with different efficacy and growth kinetics for each patient-derived model (3/6, 50% for P20-11; 5/5, 100% for P20-23; **Figures 3c-d**). Consistent with this observation, histological analyses of residual tissue post-castration highlighted the presence of a rare and small cancer foci in P20-11, while P20-23 was characterized by larger residual tumors with strong AR positivity (**Figure 3e; Figure S3)**. Thus, while both models are androgen-dependent *in vitro* and *in vivo*, the P20-11 model series appears to be more sensitive to androgen deprivation than P20-23, in line with their AR and clinical status. Moreover, the stronger sensitivity of both model series to androgen deprivation *in vivo* is in line with the higher pathway activity detected *in vivo* versus *ex vivo* (see **Figure 2**).

**Figure 3.**
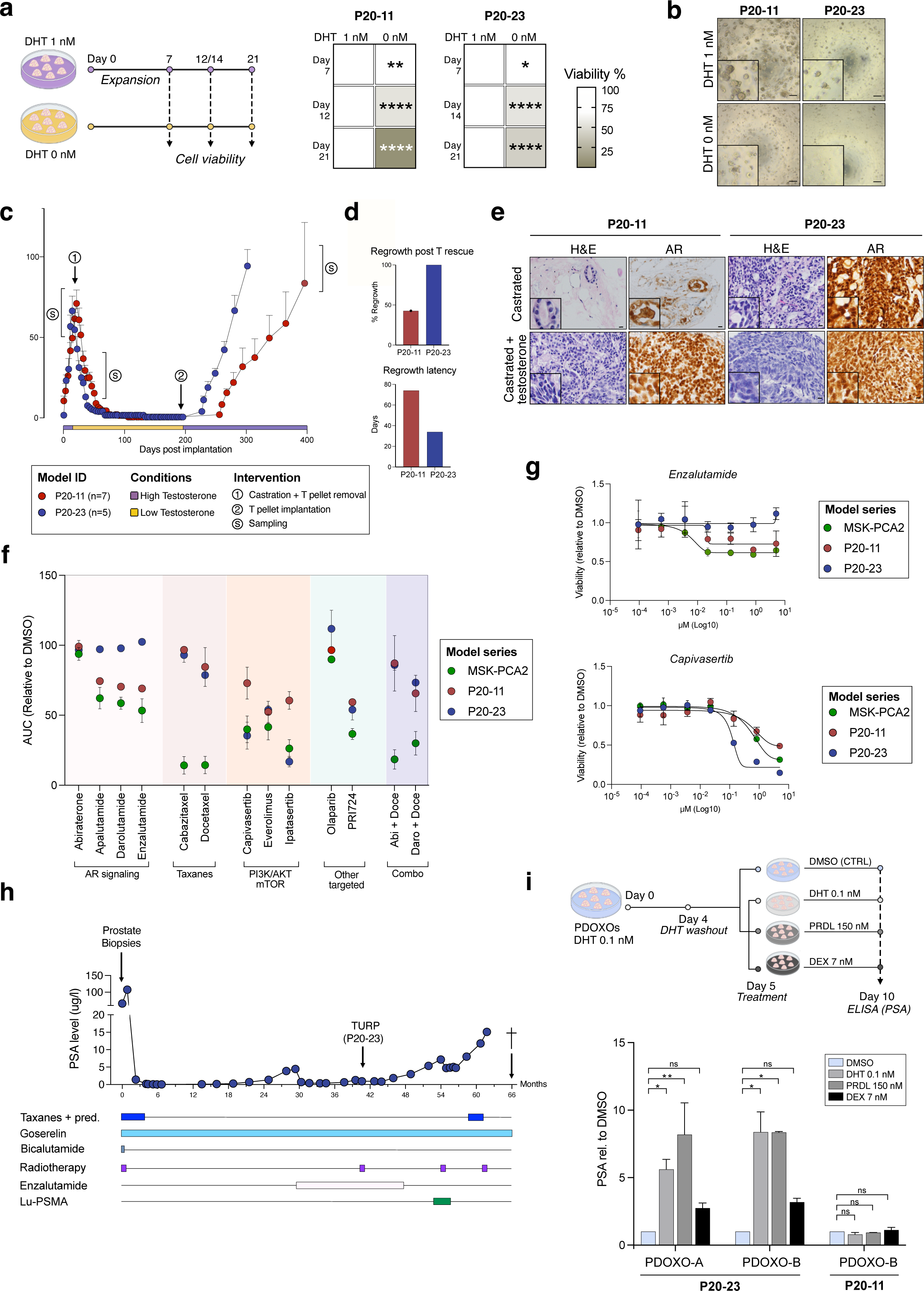
Androgen sensitivity and drug response profiles correlate with molecular and clinical attributes. **(a) Left**: Schematic representation of experimental set-up. **Right:** Viability percentage of P20-11 and P20-23 PDOXO models, relative to 1 nM for each time-point. Statistical test for P20-11 PDOXOs: One-way Anova, multiple comparisons, adjusted P values: day 7 = 0.0048, day 12 <0.0001, day 21 <0.0001. Statistical test for P20-23 PDOXOs: One-way Anova, multiple comparisons, adjusted P values: day 7 = 0.1105, day 14 < 0.0001, day 21 < 0.0001. **(b)** Brightfield images of P20-11 and P20-23 PDOXOs in medium supplemented with DHT 1 nM or DHT 0 nM after 21 days of growth. Scale bars represent 200µm. **(c)** PDOX growth kinetics of both series following castration and testosterone (T) pellet removal (1), and after testosterone pellet re-implantation (2). Colored bars on the X axis represent theoretical high and low testosterone conditions (not experimentally measured). (S) represents the timepoints of sampling **(d) Top**: Bar plot showing the percentage of mice with regrowth of tumors after testosterone pellet re-implantation. **Bottom**: Bar plot showing the latency period in days between the testosterone pellet re-implantation and the regrowth of tumors. **(e)** Representative H&E images and IHC staining for AR in the different P20-11 and P20-23 PDOX experimental groups. Scale bars for H&E and IHC images represent 20µm. **(f)** Drug profiling of P20-11 and P20-23 PDOXOs and MSK-PCA2 PDOs in Matrigel-free conditions. Drugs are grouped by mechanism of action (e.g., AR signaling) and the effect of compounds is shown as area under curve (AUC) relative to DMSO calculated from dose-response viability read-out. All treatments were performed in low androgen conditions (DHT 0.1 nM). **(g)** Dose-response curves for enzalutamide and capivasertib shown as percentage of viability relative to DMSO control. **(h)** P20-23 patient clinical and treatment course with longitudinal PSA levels starting from initial diagnostic prostate biopsies. In addition to anti-cancer treatment, denosumab was administered continuously from initial diagnosis for bone protection (not represented). **(i) Top**: Schematic representation of experimental set-up. **Bottom**: PSA levels normalized to total protein content shown as relative to DMSO (DHT 0 nM + vehicle) upon DHT 0.1 nM, prednisolone (PRDL) 150 nM or dexamethasone (DEX) 7 nM 5-day exposure and measured by ELISA in the culture medium. Statistical test for P20-23 (PDOXO-A): One-way Anova, multiple comparisons to DHT 0 nM, adjusted P values: DHT 0.1 nM = 0.0343, PRDL = 0.0077, DEX = 0.3927. Statistical test for P20-23 (PDOXO-B): One-way Anova, multiple comparisons to DHT 0 nM, adjusted P values: DHT = 0.1 nM 0.0293, PRDL = 0.0246, DEX = 0.5022. Statistical test for P20-11 (PDOX-B): One-way Anova, multiple comparisons to DHT 0 nM, adjusted P values: DHT 0.1 nM = 0.4411, PRDL = 0.9382, DEX = 0.7923. Statistical tests performed on values normalized to total protein content and non-normalized to DMSO. Replicate 1 is represented and replicate 2 can be found in figure S5.

### Organoid drug response profiles correlate with molecular and clinical attributes

The rapidly evolving therapeutic landscape for advanced PCa has made treatment selection increasingly complex. Functional precision oncology (FPO) has emerged as a promising approach to guide therapy based on the combination of molecular and *ex vivo* drug profiling (*25*); yet linking *ex vivo* results with treatment outcomes in patients, and ultimately defining the clinical relevance of such FPO strategies, remains a significant challenge. In this context, we sought to leverage our organoid models derived from non-end-stage samples and the detailed clinical follow-up of the original patients. We tested the response of P20-11 and P20-23 PDOXOs to drugs or drug combinations which were either (i) standard of care (SOC) for metastatic PCa patients (*26–28*), or (ii) tested in clinical trials for metastatic PCa patients (*7, 29*), or (iii) targeting relevant molecular pathways (**Figure 3f**). Drug testing was performed in low androgen condition (0.1 nM DHT) to mimic patients undergoing ADT (SOC) and results were compared to the well-established AR^+^ enzalutamide-sensitive PDO line MSK-PCa2 (*13*). While the CRPC-derived P20-23 organoids did not respond to any ARPis, P20-11 and MSK-PCa2 organoids exhibited some degree of response to apalutamide, darolutamide, and enzalutamide (**Figure 3f-g; Figure S4)**. None of the organoid models displayed response to abiraterone, in line with its adrenal- and testis-driven main mode of action preventing significant efficacy *in vitro*. Notably, the absence of response of P20-23 organoids to ARPis well reflected the clinical history of the patient, who was treated with enzalutamide prior to sampling and developed resistance post-TURP as evident by increasing PSA levels (**Figure 3h)**. This resistance pattern may be favored by specific *AR* genomic alterations (i.e., mutation, amplification), detected by our WES analyses in the patient tumor and organoids, and previously demonstrated to contribute to ARPi resistance (*30*).

The response to taxanes was most pronounced in the highly proliferative MSK-PCa2 organoid model and only minimal sensitivity was observed in P20-11 and P20-23 PDOXOs. The combination treatments did not provide additive effects as compared to the docetaxel monotherapy. Furthermore, all models were highly sensitive to agents targeting the PI3K/AKT/mTOR pathway (capivasertib, everolimus and ipatasertib), in line with *PTEN* alterations and pathway activity detected in the models, and highlighting the potential of such agents in targeting PTEN_null_ advanced PCa tumors. Finally, while the PARP inhibitor olaparib was inefficient, in line with absence of relevant genomic alterations, the PRI-724 β-Catenin inhibitor induced a significant response in all organoid lines, highlighting the relevance of WNT signaling in advanced PCa; nevertheless, the *CTNNB1* mutation detected in P20-11 did not appear to sensitize it further to PRI-724, possibly due to WNT pathway stimulation by compounds comprised in the medium formulation.

### Prednisolone activates the AR pathway in enzalutamide-resistant P20-23 cells

Noteworthily, the specific mutation in AR (*L702H*) detected in the P20-23 sample and derived models, was previously shown to arise upon treatment with, or post resistance to ARPis (*30*). Consistent with these studies, *AR_L702H_* was not detected in the initial diagnostic biopsy of the patient prior to any treatment (**Figure S5a)**. Among its putative function, *AR_L702H_* has been shown to be activated by certain glucocorticoids, such as prednisolone, in absence of androgens (*31, 32*). In line with this hypothesis, P20-23 organoids exhibited increased AR signaling activity upon prednisolone (PRDL) treatment, as evidenced by a significant increase in PSA secretion compared to vehicle control (8.18-fold increase in P20-23A, **Figure 3i; Figure S5b)**. In comparison, a non-significant increase was observed in both P20-23 organoids treated with dexamethasone (DEX), while no increase was detected in P20-11 organoids treated with either PRDL or DEX. Notably, P20-23-specific PRDL-driven PSA increase was observed in both PDOX-A and PDOX-B derived organoids, even though *AR_L702H_* was only detected in PDOX-A. Paralleling the results observed *in vitro*, the PSA levels of the patient rose continuously during enzalutamide treatment and further increased during subsequent treatment with cabazitaxel + prednisone (**Figure 3h**). Altogether, these data suggest that organoids mirror molecular and drug sensitivity profiles observed in matched patients and may hold predictive power.

### P20-11 PDOs enable longitudinal assessment of ADT-induced response at single-cell level

Understanding the molecular mechanisms driving PCa progression towards its most advanced stages (i.e., CRPC, NEPC) remains a long-standing challenge. Access to clinical samples after response to first-line therapy, but before relapse, remains limited; this represents a major obstacle in identifying treatment-tolerant persister cells, putatively responsible for subsequent relapses (*33*). As an alternative, the use of clinically relevant PDO models, allowing temporal assessment of drug response at single-cell resolution, represents a promising approach. In this context, we thought to leverage P20-11-derived organoids which are ADT-sensitive yet still show a small population of viable persisting cells after 3 weeks in androgen deprivation conditions (**Figure 3a-b**).

To ensure the maximum molecular overlap with the patient sample, we took advantage of our remaining P20-11 PDO vials (i.e., established directly from the patients’ sample) and performed scRNA-seq with MULTI-seq sample multiplexing (*34*) at two time points and in the presence or absence of androgens (**Figure 4a**). While no transcriptomic shift between the two conditions was observed after 3 days (**Figure 4b, top panel**), a clear separation occurred after 21 days in culture (**Figure 4b, bottom panel**), suggesting major transcriptomic reprograming. A key feature of this transcriptomic dynamic was well illustrated by the *ANDROGEN RESPONSE (*MSigDB, Hallmark) signature score, which remained stable after three days but showed a pronounced decline in the androgen-deprived condition after 21 days (**Figure S6a)**.

**Figure 4.**
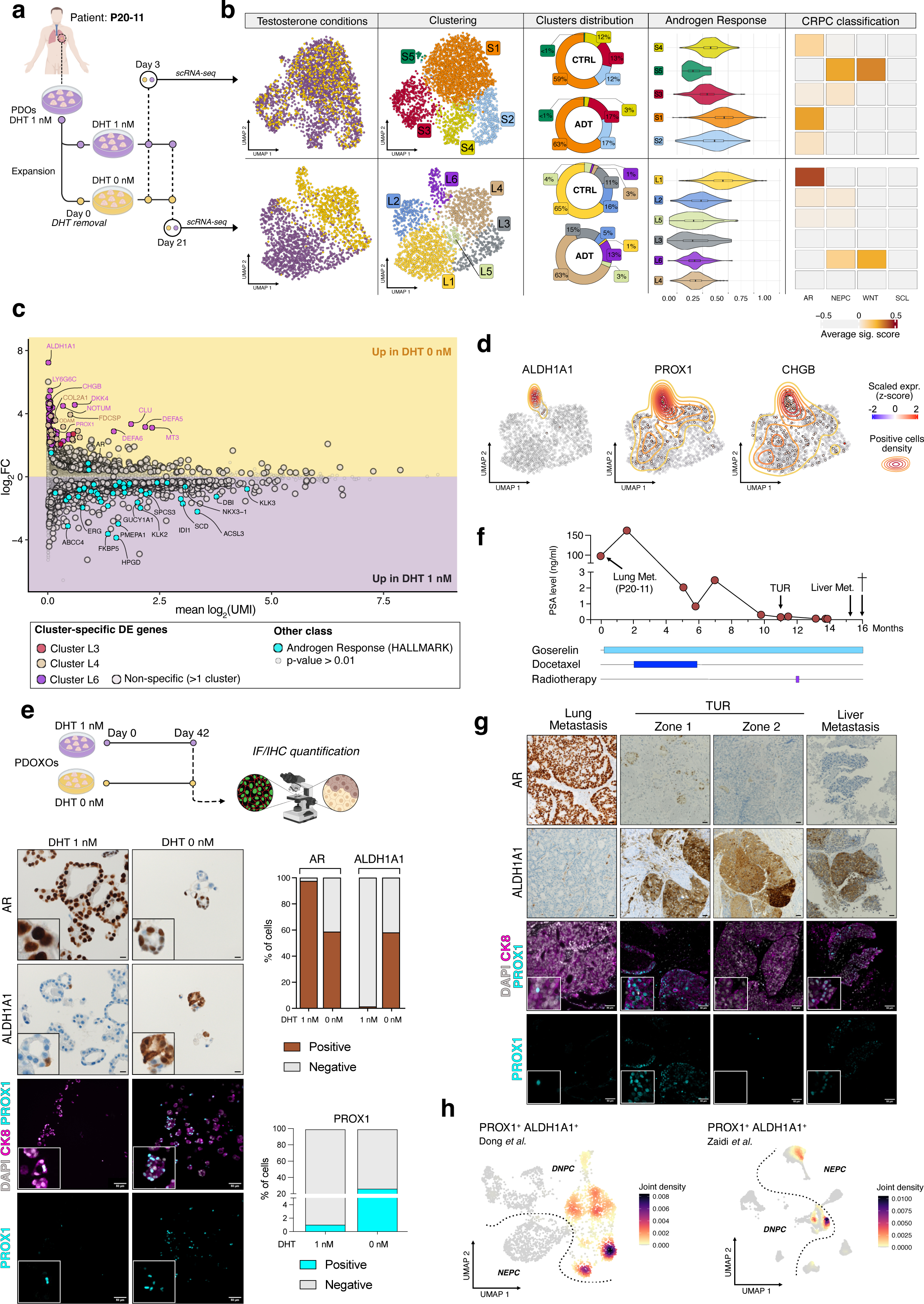
Longitudinal scRNA-seq analysis of P20-11 organoids upon ADT uncover molecular traits consistent with the disease progression path of the original patient. **(a)** Schematic representation of experimental set-up. **(b)** Clustering and *ANDROGEN RESPONSE* signatures (MsigDB, Hallmark) of P20-11 PDOs in control (DHT 1 nM) and androgen deprivation conditions (DHT 0 nM). **Top Row**: day 3. **Bottom row**: day 21. **From left to right**: UMAP projection of control and androgen-deprived PDOs at day 3 versus day 21. Unsupervised clustering results from PDOs (day 3 clusters: S1 – S5, day 21 clusters: L1-L6). Cluster distribution in control and treated conditions, highlighting the relative abundance of each cluster in each condition. Violin plot of *ANDROGEN RESPONSE* signature score (MsigDB, Hallmark) shown by cluster. Heatmap visualization of signature scores per cluster using previously published CRPC sub-type signatures (*35*). **(c)** MA plot of differentially expressed genes in PDOs cultured in DHT control conditions (1 nM) or in DHT-deprived conditions (0 nM) at day 21. Genes associated with the *ANDROGEN RESPONSE* signature score (MsigDB, Hallmark) are highlighted in blue. Genes specifically upregulated in cluster L3, L4 and L6 are also highlighted. **(d)** UMAP projection of day-21 cells with density overlays of gene-positive populations. Cells are color-coded according to their scaled expression, and density overlay of cells with a positive scaled expressed is shown as red lines. **(e) Top**: Schematic representation of experimental set-up. **Bottom left**: Immunohistochemistry analysis with antibodies recognizing AR and ALDHA1 as well as immunofluorescence analysis with antibodies recognizing CK8 and PROX1 in PDOXOs cultured in androgen proficient (DHT 1 nM) and androgen deficient (DHT 0 nM) conditions. **Bottom right**: Bar plots showing the percentage of cells positive for AR, ALDH1A1, and PROX1 cultured in DHT 1 nM and DHT 0 nM, as quantified from immunohistochemistry (AR and ALDH1A1) and immunofluorescence analyses (PROX1). Scale bars represent 20µM for IHC images and 50µM for IF images. **(f)** P20-11 patient clinical and treatment course with longitudinal PSA levels starting from initial diagnostic lung metastasis. **(g)** Phenotypic characterization of P20-11 patient samples at initial diagnostic (lung metastasis) and following progression samples (TUR and liver metastasis). Shown are representative IHC and immunofluorescence staining for the indicated antibodies. Scale bars represent 20 µM for IHC images and 50 µM for IF image. **(h)** UMAP projection of double-negative prostate cancer (DNPC) and neuroendocrine prostate cancer (NEPC) cells from single-cell RNA-sequencing cohorts. Data from Dong et al. (*41*) (left) and Zaidi et al. (*42*) (right) are shown. Dotted lines indicate the boundary between DNPC and NEPC states. The density of PROX1/ALDH1A1 double-positive cells is color-encoded, highlighting this subpopulation across both entities and datasets. Representative single-marker expression and sample distributions are shown in Supplementary Figure 8.

Optimized clustering analysis identified five cell clusters at day 3 (short-term: clusters S1-5) and six cell clusters at day 21 (long-term: clusters L1-6), with cell cycle contributing partially to the separation of the subpopulations (**Figure S6b-c**). Notably, cell cluster distributions were largely similar between conditions at day 3, except for cluster S4. This cluster was identified by GSEA as an OXPHOS/translation-active cluster (**Figure S6d; Dataset S1**) and accounted for only 3% of cells in the ADT condition versus 12% in the control, suggesting that ADT induces an early reduction in metabolic activity. In contrast, by day 21, treatment had markedly altered the cellular cluster composition. In the control group, most cells (65%) belonged to cluster L1, characterized by upregulated AR signaling and lipid metabolism, (**Figure S6e**), similarly to cluster S1. However, in the ADT group, the majority of the cells (63%) was assigned to cluster L4, displaying reduced AR signaling and expressing stress/apoptosis-related genes such as the cell cycle arrest cyclin CCNG2. Proliferative cells (cluster L2) were less present under androgen deprivation (5% vs 16%). Clusters L3 and L5 were evenly distributed between treatment conditions and enriched for OXPHOS/translation terms, likely deriving from cells grouped under cluster S4 at day 3. Finally, cluster L6 was enriched for WNT signaling and epithelial–mesenchymal transition pathways. Differential expression analysis indicated that these cells share the same top genes than the cells of cluster S5 at day 3 (**Figure S6f-g**; **Dataset S1**). Importantly, while cells in cluster S5 were present in both conditions at the same ratio (< 1%), cluster L6 showed a relative increase after 21 days of androgen deprivation (1% to 13%). Notably, clusters S5 and L6 displayed low androgen-response (HALLMARK) scores and were assigned to the AR-negative NEPC-CRPC and WNT-CRPC subtypes, based on recently-defined CRPC transcriptomic signatures (*35*) (**Figure 4b**).

### Androgen deprivation enriches in AR-negative ALDH1A1^+^ PROX1^+^ castration-tolerant persister cell populations

In an effort to narrow down transcriptomic changes associated with ADT and identify putative castration-tolerant persister cell populations, we performed DEA between both treatment groups at day 21 (**Figure 4c; Dataset S1**). Strikingly, expression of a subset of AR-regulated genes (e.g., FKBP5) became nearly undetectable in the ADT group, while others such as KLK3 and KLK2 were reduced to a lesser extent (**Figure S6h**). We then pinpointed top treatment-induced genes driven by single clusters, mostly focusing on ADT-enriched cluster L4 and L6. Notably, cluster L6–specific genes displayed some of the most enriched genes under ADT. These included ALDH1A1, a marker previously linked to the WNT/beta catenin pathway, stemness and PCa progression (*36, 37*), the neuroendocrine-associated gene CHGB, and PROX1, a recently-identified driver of lineage plasticity in PCa (*38–40*) (**Figure 4d).** Notably, PROX1 expression was already detectable at day 3 in cluster S5, reinforcing its putative link with cluster L6 at day 21. In contrast, ALDH1A1 and CHGB showed either no or only background expression in cluster S5, indicating that these transcriptional programs are likely acquired at later stages (**Figure S6i**).

To validate these findings at protein level and within timeframes better mirroring sustained long-term ADT, we leveraged PDOXOs cultured in androgen-deprived or -proficient conditions for 42 days (**Figure 4e**). In line with our scRNA-seq data in PDOs, cells positive for ALDH1A1 and/or PROX1 were present at very low percentages in androgen-proficient organoids but significantly increased in androgen-deficient conditions (1.37 versus 58.65 % ALDH1A1^+^ and 1.07 versus 26.06% PROX1^+^ cells; **Figure 4e; Figure S7a, Table S2**). These cell dynamics were accompanied by marked reduction of AR+ cells in androgen-deficient organoids; despite expression of the early driver of lineage plasticity PROX1, cell positive for terminally-differentiated neuroendocrine markers, such as CHGA and Synaptophysin, were not clearly detected in either condition (**Figure S7b**). Taken together, these data suggest that ADT selects for cells expressing early molecular programs of stemness and neuroendocrine lineage in organoids.

### Organoids uncover molecular traits consistent with the disease progression path of the original patient

In light of these findings, we sought to re-analyze samples obtained from the patient at initial and later stages of his disease course; samples included the untreated lung metastasis specimen from which the P20-11 model series was derived from, as well as post-treatment local progression (transurethral resection: TUR) and liver metastasis samples (**Figure 4f**). Similar to control organoids, the original treatment-naïve sample was characterized by a large majority of tumor cells expressing AR, and very rare cells positive for ALDH1A1 and/or PROX1 (**Figure 4g**). In stark contrast, the TUR progression sample was largely comprised by AR-negative tumor cells as well as rare focal AR-low cells while the liver metastasis was exclusively composed of AR-negative tumor cells, consistent with PSA level dynamics detected in the patient (**Figure 4f**). Furthermore, both post-progression samples comprised tumor areas with ALDH1A1 and/or PROX1 positivity, reminiscing castration-tolerant phenotypes observed in organoids. Rare areas of focal neuroendocrine differentiation were also found in the TUR sample (**Figure S7c**). To broaden our findings, we re-analyzed recently published scRNA-seq cohorts, which highlighted an enrichment of PROX1 and ALDH1A1 expression in a subset of AR-negative/low CRPC patient samples (*41, 42*) (**Figure 4h**, **Figure S8**). Altogether, these analyses demonstrate that organoids mirroring the biology of tumors provide clinically relevant models for inferring disease progression and identifying pertinent biomarkers in PCa.

## Discussion

The establishment of long-term expandable organoid lines derived from advanced PCa clinical specimens still remains a significant challenge in the PCa research field, as reflected by the limited number of sharable models (*13, 17, 35, 43*). While the xenotransplantation of promising PCa PDO cultures allows demonstrating their tumorigenicity capacity *in vivo* and expanding the material, it remains particularly inefficient in terms of take-rate and speed (*16*). This is well illustrated by the long latency period for tumor initiation in our xenograft models generated from PDOs. Furthermore, we show that subpopulations of cells with clinically relevant genomic features may be selected in/out *in vivo* and *in vitro*, highlighting the need for thorough molecular characterization of models through different generations, sublines, and model types. On the other hand, molecular differences in sublines may provide unique opportunities to assess the relevance of specific mutations in an isogenic setting.

With the strengthening of the 3Rs (Replace, Reduce, Refine) principles and the recent FDA plans to phase out animal testing requirement for preclinical drug testing, organoid-based models are becoming increasingly important in translational cancer research. In this context, xenograft models are frequently re-cultured as organoids in various cancer types including PCa (*15, 44–48*), allowing for reduction of animal experimentation and for larger-scale experiments. While our single-cell level comparisons of PDOX and PDOXO models reveal some differences at gene-level, we show that the core biologically relevant pathways remain conserved, supporting the stability and relevance of organoid systems. Taking in account our restricted pairs of PDOXs/PDOXOs, we encourage others to pursue similar comparisons to build trust in organoids and identify pertinent and irrelevant applications for these models. In this framework, it is worth highlighting that outcomes of drug testing with certain compounds should be considered with caution when tested in PDOXOs/PDOs. As an example, abiraterone is typically not effective *in vitro* and taxane responses may depend on organoid doubling times due to their microtubule-disrupting mechanism. This is illustrated by our own analyses, with the MSK-PCa2 PDO line proliferating at a substantially higher rate and exhibiting significantly higher sensitivity to cabazitaxel and docetaxel, as compared to P20-11 and P20-23 PDOXOs. Finally, it remains to be demonstrated whether the sensitivity of all organoid models to PI3K/AKT pathway inhibitors is directly linked to their molecular characteristics (i.e., genomic alterations in *PTEN*) and/or is a consequence of pathway-activating factors comprised in the culture medium.

Beyond these technical considerations, our work highlights the urgent need for validating the relevance of PDOs in prospective patient/organoid co-clinical studies (*49*). Given that most PDOs are generated from patients with end-stage diseases through autopsy programs or from patients who do not undergo systemic treatment, direct patient/organoid comparative studies are currently lacking. Here, as an alternative, we retrospectively show that P20-23 organoids and the original patient are both enzalutamide resistant and carry *AR* alterations which have previously been demonstrated to drive ARPi resistance (*50*). Furthermore, we demonstrate that prednisolone (the active form of the prodrug prednisone used in the patient) activates the AR pathway in P20-23 PDOXOs, consistent with prior studies linking GR signaling to resistance to androgen-targeting therapy (*31, 50, 51*). Notably, prednisolone-mediated AR activation specifically occurred in P20-23 organoids, irrespectively of the presence of *AR_L702H_*, suggesting an effect intrinsic to P20-23 tumor cells which is (at least partially) independent from a promiscuous *AR* mutation. Complementing previous works linking molecular and drug response features (*15, 44*), our comparative organoid/patient data provide a proof-of-concept for the relevance of organoid-based FPO strategies in PCa; while we acknowledge their correlative nature, our analyses suggest that the original patient may have not benefited from standard-of-care treatment regimens he received (i.e., enzalutamide, taxanes + prednisone), and highlight the utility of genomic sequencing before decision making in specific advanced PCa cases. Of note, our approach of leveraging organoids to predict multiple drug sensitivities in dose-response assays would not have been possible without prior expansion *in vivo*; this further highlights the challenge of such endeavor for personalized medicine applications which require quick clinical decision making.

Furthermore, the newly-generated P20-11 model series provides a rare opportunity to study the transition between the HSPC and CRPC states given that (i) it originates from an hormone-naïve metastatic specimen with wild-type *AR,* a disease state with few available experimental models (*52, 53*), and (ii) exhibits genomic alterations (*TP53*, *PTEN*, *RB1*, *CTNNB1*) which are associated with lineage plasticity and aggressive CRPC subtypes such as NEPC and DNPC (*54–56*). Exploiting this model, our scRNA-seq longitudinal analyses uncovered ADT-tolerant cell populations, associated with AR-negative subtypes and lineage plasticity transcriptomic features. In spite of these features priming the cells towards NEPC differentiation, we could not detect cells expressing classical neuroendocrine markers at protein level; noteworthily, transdifferentiation from adenocarcinoma towards NEPC has been achieved in established cell lines and PDXs (*57*) but has not yet been demonstrated using human PCa organoids *in vitro*, suggesting that current PDO culture conditions may not be prone to such cellular transformation. Finally, re-growth of the PDOXs models post-castration only occurred upon supplementation of testosterone, highlighting their androgen-dependence, irrespective of their origin (CRPC or HSPC patients). This emphasizes the distinction between patient clinical state and experimental model biology, particularly in the context of androgen sensitivity which may be highly dependent on the *AR* status and baseline experimental conditions (i.e., testosterone levels *in vivo*, DHT concentrations *in vitro*) (*53*).

To the best of our knowledge, our longitudinal single-cell data of P20-11 PDOs provide first evidences that PDOs can mirror patient-specific molecular and clinical trajectories in the context of PCa. Our analyses uncover PROX1, a factor recently linked to NE differentiation and liver metastasis (*38–40*), and reinforce its importance in a clinically relevant context of matched patient- and model-longitudinal samples. Furthermore, our data bring re-emphasize ALDH1A1 as a marker for castration-tolerant cells (*58*), shedding new light on its potential relevance for progression towards AR-negative subtypes. Finally, our analyses uncover a subset of cells surviving androgen deprivation which, although rare, pre-exist in the treatment-naïve setting, supporting the hypothesis of the pre-existence of castration-resistant populations in PCa patients (*59–61*). In summary, P20-11 and P20-23 model series can be expanded *in vitro/in vivo* and exhibit molecular and functional features that are representative of advanced PCa. Through organoid/patient comparative analyses, we provide initial evidences supporting the relevance of PCa organoid models for FPO strategies, mechanistic insights, and biomarker identification.

## Material and Methods

### Patients and clinical samples

The two parental patients’ samples were obtained as previously described in Servant et al (*14*) under approval by the Ethics Committee of Northwestern and Central Switzerland (EKBB 37/13). The P20-11 sample was obtained from a resected specimen of a hormone naïve prostate cancer metastasis in the lung. The patient had a PSA of 97,80 μg/l at diagnosis, was treated continuously with ADT (goserelin), and received 6 cycles of chemotherapy (docetaxel) with excellent response (PSA <1 μg/l). The patient progressed 5 months later and underwent a palliative TUR. A liver metastasis was detected 5 months later as the disease continued to progress. The patient passed away 1 year after the initial diagnosis.

The P20-23 sample was obtained from a castration-resistant transurethral resection of the prostate (TURP). The patient was originally diagnosed with metastatic PCa, ISUP Grade Group 5; ADT (goserelin) was started as well as docetaxel complemented with prednisone. Docetaxel was interrupted after 6 months. Enzalutamide was added to ADT 30 months after the initial diagnosis as the cancer was progressing. The TURP from which PDOs were derived was performed 9 months after starting enzalutamide (PSA: 1,27 μg/l, ISUP5). The patient next received 6 cycles of Lu_177_-PSMA, followed by 4 cycles of chemotherapy (cabazitaxel + prednisone), entered palliative care, and passed away 5 years after the initial PCa diagnosis. Detailed treatment timeline and clinical courses of the patients are represented in **Figure 3h** and **Figure 4f**.

### Xenograft and organoids models

#### Generation and passaging of patient-derived organoids xenografts (PDOXs)

The two corresponding PDO lines were maintained in Matrigel domes for 18 days (P20-11) and 29 days (P20-23), as described in Servant et al (*14*). Single-cell suspensions from both PDO lines were generated prior to *in vivo* injection for PDOX establishment. To achieve this, the culture medium was replaced with Dispase (1 mg/mL) and the Matrigel domes were gently disrupted. After a 30min incubation at 37 °C, organoids were collected and resuspended in TrypLE Express. Complete single-cell suspension was achieved through successive 5-min digestion rounds, with microscopic inspection between each. Optimal digestion time was reached within 5-10 min and the reaction was quenched by adding an excess of cold adDMEM/F12^+++^. Cells were then centrifuged and viability was assessed using an automated cell counter (Countess II FL, Thermo Fisher Scientific).

Cells were resuspended in 50:50 Matrigel:PBS, and injected subcutaneously in the flank of male NOD Scid Samma (NSG) immunodeficient mice, aged 6 to 8 weeks old. All mouse experiments were conducted with the approval of the Animal Care Committee of the Kanton Base-Stadt, Switzerland (approved licenses #3053, #3066-32428, and #3066-35110). Mice were bred and maintained in the animal facility of the Department of Biomedicine of the University of Basel under specific pathogenic-free conditions on a 12h day and 12h night schedule with ab libitum access to food and drinking water. For P20-11 PDOX, a testosterone pellet was implanted in mice 2 days before engraftment to increase testosterone levels of the host. Mice were monitored bi-weekly and sacrificed when the endpoint was reached, as per stated in the ethical authorizations. To establish the first generation of PDOXs, 120 000, and 67 500 cells were injected to generate P20-11 PDOX-A and P20-11 PDOX-B, respectively (tumor growth in 2/6 mice); 200 000 cells per mouse were injected to generate P20-23 PDOX-A/B/C (tumor growth in 3/3 mice). Additional details regarding PDOX models used for analyses can be found in **Table S1**.

#### Xenograft processing, passaging, and xenograft-derived organoid culture

After sacrifice and tumor collection, PDX tissues were cut into ∼1 mm³ pieces using scissors and a scalpel. Tissue fragments were incubated in a pre-warmed digestion mix containing collagenase type IV (2 mg/mL), Dispase I (1 mg/mL), and DNase I (100 µg/mL) in adDMEM/F12^+++^ supplemented with ROCK inhibitor (10 µM). Digestion was carried out in Petri dishes at 37°C for 30–60min, with gentle agitation and intermittent pipetting to help release the cells. The cell suspension was then rinsed with adDMEM/F12^+++^ (5 min, 300 × g), and TrypLE Express supplemented with ROCK inhibitor and DNase I was used to complete tissue dissociation. Optimal digestion was typically achieved within 10–15 min and did not exceed 20 min to preserve cell viability. The reaction was quenched by adding an excess of cold adDMEM/F12^+++^ and filtered through 100 µm strainers to obtain a clean single-cell suspension. Cells were centrifuged, counted by Trypan Blue exclusion, and used either for re-injection into mice (PDX passage), organoid generation, or downstream applications such as single-cell RNA sequencing (scRNA-seq). When required, additional steps such as red blood cell lysis, finer filtration, or dead-cell removal were performed to optimize viability and purity.

For xenograft passaging, 1million of live cells were re-injected into the flank of NSG mice. For organoid generation, cells were resuspended in a mixture of 75% Matrigel and 25% adDMEM/F12 at a density of 500–1,000 cells/µL, and 13–25 µL drops were plated per well depending on the plate format. Plates were incubated upside down at 37°C for 30 min to allow Matrigel polymerization before adding the appropriate volume of culture medium containing ROCK inhibitor and DHT according to their clinical status (i.e., 1 nM DHT for P20-11, 0.1 nM DHT for P20-23).

### *In vivo* castration experiments

All experiments involving animals were conducted in accordance with a protocol approved by the Swiss Veterinary Authority/Board (TI-70-2024) and received approval from the ethical committee of the Institute of Oncology Research (IOR). *In vivo* castration experiments were performed using the P20-23A and P20-11B models. Xenograft tumors were established in 6–8-week-old NOD-Rag1^null^ IL2rg^null^ (NRG) immunodeficient mice through subcutaneous injection of single-cell suspensions derived from dissociated PDX tissue, as previously described. For the 20-11 model, a testosterone pellet was implanted one week prior to injection to support tumor engraftment. Mice were weighed every two weeks before tumors became palpable and weekly thereafter. Once palpable, tumor growth was monitored twice weekly by caliper measurements. When tumors reached a volume of 50–100 mm³, a subset of mice was sacrificed for tissue characterization (intact group). The remaining animals underwent surgical castration once tumors reached 50–75 mm³, with concurrent removal of the testosterone pellet. Three weeks post-castration, a group of mice was euthanized during the regression phase, and residual tumors were collected for analysis (castrated group). The remaining cohort was maintained to monitor potential tumor regrowth. As no regrowth was observed in either model, a high-dose testosterone pellet was re-implanted 180 days after the initial xenograft, and tumors that regrew were collected for phenotypic characterization (regrowth group).

### *In vitro* androgen deprivation and drug testing

After dissociation of the PDOXOs to a single cell suspension, 5000 single cells per well were plated in 96well/plates, in 10 μL domes with 75% GFRM. Organoids were grown in organoid media containing 1 nM or 0 nM of DHT from day 0 after plating until the end of the experiment.

For drug testing experiments, 1000-2500 PDOXO-derived single cells per well were plated on ice in ultra-low attachment 384-well plates (Phenoplate #6057300) and organoid medium. Organoids were allowed to grow for 3 to 5 days before treatment was started. Organoids were treated for 5 days with vehicle control (DMSO 0.1%) or drugs diluted in DMSO. Drugs and DMSO were dispensed with the Tecan D300e using the randomization feature. The drugs used were enzalutamide (1 nM – 30 μM), abiraterone (1 nM – 50 μM), apalutamide (0.1 nM – 10 μM), docetaxel (0.01 nM – 1 μM), PRI-724 (1.6 nM – 8 μM), cabazitaxel (0.01 nM – 1 μM), darolutamide (1.3 nM – 10 μM), capivasertib (0.1 nM – 5 μM), everolimus (1.3 nM – 10 μM), ipatasertib (0.8 nM – 20 μM), olaparib (2.8 nM – 30 μM), all purchased from SelleckChem. Cell viability was assessed with the CellTiterGlo3D luminescence assay (Promega). All drug treatment experiments were conducted in triplicates and two biological replicates were performed. Dose response analysis and Area Under the Curve (AUC) were assessed using GraphPad Prism9.

### Histology, immunohistochemistry, and immunofluorescence

Histological analysis of the samples was performed with standard hematoxylin and eosin (H&E) staining. ISUP grading and histological characteristics of the tissues were defined by a genitourinary pathologist. Immunohistochemistry (IHC) and Immunofluorescence (IF) stainings were performed as previously described (*14*). H&E and IHC images were captured with a 40x objective, unless stated otherwise in the legend of the figure. IHC/IF image quantification was performed using QuPath 0.4.3. Cells were segmented using the Cell Detection plug-in followed by the Single Measurement Classifier to classify cells as positive or negative, based on a user-defined threshold (**Table S2**). Antibodies and dilutions are provided in the **Table S3**.

### Prostate Specific Antigen (PSA) ELISA (*in vitro*)

The concentration of PSA secreted in the culture media by the organoids at the end of the experiment was measured via ELISA (ref 0208, Abnova), according to manufacturer’s protocol. Culture media was stored at −20°C before performing the ELISA and was diluted 1:2 with the Zero Buffer to ensure that all samples fit within the standard curve and minimize the matrix effect. Absorbance 450nm was read with the VarioSkan Lux (Thermofischer). Data was analyzed using GraphPad Prism 9 and one-way ANOVA. The concentration of PSA/ml was standardized to the total amount of protein in the sample, measured with a Bradford assay according to the manufacturer’s protocol (Bio-Rad Protein Assay). The results are expressed as amount of PSA (ng) per mg of total protein in the culture media and normalized to vehicle control (DHT 0 nM).

### DNA extraction

Patient tissues were stored at −80°C and crushed to a powder in liquid nitrogen prior to extraction. Dispase (1mg/ml) was used to extract the organoids from the Matrigel. After a PBS wash, organoid pellets were frozen and stored at 80°C until extraction. Germline control was obtained by extracting DNA from Peripheral Blood Mononuclear Cells (PBMC) of the patients. PBMC were separated from other blood cells using Ficoll and PBMC pellets were stored at −80°C until DNA extraction. The *Quick*-DNA/RNA Miniprep Plus kit (Zymo Research) was used to extract DNA from blood, tissues and organoids, as per guidelines provided in the kit. DNA concentrations were measured using QuBit high sensitivity kit (Invitrogen).

### Whole exome sequencing and variant annotation

Extracted DNA from flash frozen tissue samples and their matched germline components were subjected to WES. Twist Human Core Exome + RefSeq + Mito-Panel kit (Twist Bioscience) was used for the whole exome capturing according to manufacturer’s guidelines. Paired-end 100-bp reads were generated on the Illumina NovaSeq 6000. After the sequencing, reads were aligned against the hybrid reference human/mouse genome GRCh38/GRCm38, as previously described using Burrows-Wheeler Aligner (BWA, v0.7.12)(*62*). Single Nucleotide variants (SNVs) and indels were called using MuTect2 (GATK 4.1.4.1) (*63*). SNVs or indels with a VAF < 1% or that were covered by fewer than three reads were discarded, if the SNVs were found in both components of one sample, a cut-off of two reads was applied. We further excluded variants identified in at least two of a panel of 210 non-tumor samples, including the non-tumor samples included in the current study. Variant annotation was performed by SnpEff software v.4.1 (*64*). The heatmap of non-synonymous mutations and CNV was generated using the R package MAFTOOLS v.2.0.16 (*65*) by selecting the prostate cancer driver genes from the Intogen database (*66*) and Bailey et al (*67*).

### Copy number variations (CNV)

Allele-specific CNVs were identified using FACETS v.0.5.14 (*68*) and the log ratio relative to ploidy was used to call deletions, losses, gains, and amplifications. Genes with total copy number greater than gene-level median ploidy were considered gains; greater than ploidy + 4, amplifications; less than ploidy, losses; and total copy number of 0, homozygous deletions. Somatic mutations associated with the loss of the wild-type allele (i.e., loss of heterozygosity (LOH)) were identified as those where the lesser (minor) copy number state at the locus was 0 (*69*). For chromosome X, the log ratio relative to ploidy was used to call deletions, loss, gains, and amplifications (*70*). All mutations on chromosome X in male patients were considered to be associated with LOH. CNV similarities between different samples was calculated using a Pearson’s correlation method implemented in the Corrplt package (v.0.95).

### Single-cell RNA-sequencing

Single-cell RNA sequencing (scRNA-seq) was performed on single-cell suspensions derived from xenograft tissues (PDOXs) and corresponding organoid models (PDOs and PDOXOs). Preparation of single-cell suspensions from PDOX tumor tissues followed the same dissociation protocol used for generating PDOXOs from xenografts described above. Samples showing a viability below 80% were processed with a magnetic dead cell removal kit (Miltenyi Biotec) according to the manufacturer protocol to enrich for live cells.

### Multi-Seq multiplexing method

When applicable - such as for parallel processing of multiple PDOX or organoid samples, or in drug-treatment experiments - we employed the MULTI-seq multiplexing method for single-cell RNA sequencing (scRNA-seq), following the protocol described by McGinnis et al. (*34*). This technique uses lipid-modified oligonucleotides (LMOs) to tag cells in suspension, allowing multiple pooled samples to be processed as a single reaction, thereby minimizing technical bias, and reducing costs. Briefly, following generation of single-cell suspensions, cells from each sample were sequentially labeled with an anchor LMO, a co-anchor LMO, and a unique DNA barcode. To prevent potential mislabeling caused by residual unbound LMOs, non-anchored LMOs were quenched by adding 1% BSA and removed by centrifugation at 300 × g for 5 min prior to pooling the samples. After pooling, the barcoded samples were processed as standard single-cell suspensions for library preparation and sequencing. Sample identities were later resolved bioinformatically based on their unique MULTI-seq barcode sequences.

### Library preparation

Single-cell gene expression libraries were prepared according to the Chromium Next GEM Single Cell 3’ v3.1 Dual Index User Guide (10x Genomics, Revision B). Briefly, samples meeting viability and cell number criteria were resuspended at a concentration of 700–1,200 cells/µL and loaded into the Chromium Next GEM Chip G to reach a recovery of 10,000 cells per sample. Immediately following GEM generation, reverse transcription of captured mRNA was performed to incorporate barcoded primers, including Unique Molecular Identifiers (UMIs) and 10x GEM Barcodes to the captured mRNA. Following Post-GEM-RT cleanup, PCR amplification of the first-strand cDNA was performed by PCR (11 cycles), followed by SPRIselect bead-based size selection (Beckman Coulter, B23317) to enrich for cDNA fragments of the desired size. Enzymatic fragmentation, end repair, A-tailing and ligation steps were performed immediately after. Finally, libraries were constructed by adding Illumina P5/P7 adapters and i5/i7 sample indexes via Sample Index PCR (9-10 cycles).

### Sequencing

All generated libraries were evaluated for quantity and quality using a Thermo Fisher Scientific Qubit 4.0 fluorometer with the Qubit dsDNA HS Assay Kit (Thermo Fisher Scientific, Q32851) and an Advanced Analytical Fragment Analyzer System using a Fragment Analyzer NGS Fragment Kit (Agilent, DNF-473), respectively. The libraries were pooled with other 10 x Genomics libraries and sequenced with a loading concentration of 300 pM, symmetrical paired-end and dual indexed, using various illumina NovaSeq 6000 Reagent Kits v1.5 (100 or 200 cycles), on an Illumina NovaSeq 6000 instrument. The read set-up was as follows: read 1: 29 cycles, i7 index: 10 cycles, i5: 10 cycles and read 2: 89-91cycles. The quality of the sequencing runs was assessed using illumina Sequencing Analysis Viewer (illumina version 2.4.7) and all base call files were demultiplexed and converted into FASTQ files using illumina bcl2fastq conversion software v2.20. For each sample, sequencing depth was targeted at approximately 40,000 reads per cell for endogenous RNA and 5,000 reads per cell for MULTI-seq barcode reads, as recommended in the original MULTI-seq publication (*34*). All steps post library preparation were performed at the Next Generation Sequencing Platform, University of Bern.

### Alignment and demultiplexing

Sequencing data were mapped to the hg38 genome using the STARsolo framework (v2.7.10a) and gene annotation was performed with a custom GTF file (Ensembl 104), filtered to include only transcripts of biotypes “protein_coding,” “lncRNA,” and all B- and T-cell receptor gene transcripts. The cell barcode whitelist for STAR was provided by 10x (file: 737K-august-2016.txt). Non-standard command line options for STAR included parameters for filtering, mapping, output formatting, and handling cell barcodes and UMIs, such as “–outFilteType BySJout,” “–outFilterMultimapNmax 10,” “–outSAMtype BAM SortedByCoordinate,” and various options specific to the handling of cell barcodes (CB) and unique molecular identifiers (UMIs). All subsequent data processing was conducted in R using Bioconductor (R version 4.2.2, Bioconductor version 3.16).

### Empty droplet detection, LMO demultiplexing and quality controls

Empty droplets were filtered out using the emptyDrops(niters = 50,000) function from the DropletUtils package (v1.18.1). LMO demultiplexing was performed using the package deMULTIplex2 (v 1.0.2) with default parameters. Cells with fewer than 10,000 total reads, fewer than 1,000 detected genes, more than 25% mitochondrial content, or those identified as doublets by scDblFinder (expected doublet rate = 0.076) were excluded from further analysis.

### Sample integration

To compare transcriptomic similarities and differences between PDOX and PDOXO models across passages in both P20.11 and P20.23 lineages, all single-cell experiment (SCE) objects were first rescaled to account for differences in sequencing depth between batches using the MultiBatchNorm(min.mean = 0.1) function from the Batchelor R package (v1.14.1). Variance modelling of gene expression was then performed independently for each sample using the modelGeneVar() function from scran (v1.26.2); results were combined with combineVar(), and the top 1,000 most Batch correction was considered unnecessary based on the visual inspection of sample placement on the UMAP, which showed no batch-driven segregation. highly variable genes were selected using getTopHVGs() for subsequent analyses.

### Clustering analysis

Graph-based clustering (Louvain algorithm) was performed on PCA-reduced data across a grid of parameters varying the number of nearest neighbors (k = 15–90), resolution (0.4–1.0), and number of PCs (10–100). Cluster robustness was evaluated through 50 bootstrap iterations (80% subsampling), and stability was quantified using the Jaccard index between original and resampled cluster assignments. The optimal parameter set choice was evaluated using the package scclusteval (v1.0), as the one for which the number of stable clusters most closely matched the total number of clusters, reflecting optimal agreement between cluster resolution and reproducibility.

### Differential gene expression analysis

To identify differentially expressed (DE) genes across cell groups, the FindAllMarkers() function from the Seurat package (v5.0.1) was employed using the Wilcoxon test. The results were visualized as heatmaps using the ComplexHeatmap package (v2.14.0), with genes sorted by ascending adjusted P value.

### Signature score calculation

After clustering analysis, clusters were characterized by identifying enrichment for particular gene signatures. Gene signatures were retrieved from the MSigDB database with the help of the R package msigdbr (v7.5.1) and the function AddModuleScore() built within Seurat (v5.0.1) was then used to generate a signature score for each cell.

### Pathway activation score

Pathway variability was analyzed using Pagoda2 on raw count matrices (log-scaled). Gene-wise variance was adjusted (adjustVariance, gam.k = 10), PCA computed on the top 1,000 overdispersed genes (nPcs = 20), and UMAPs generated (n_neighbors = 300, min_dist = 0.7). Pathway overdispersion was tested on MSigDB HALLMARK gene sets (30–200 genes; correlation threshold = 0.9), and pathways were ranked by dispersion scores. For each top pathway, the first principal component (PC1) projection per cell was extracted and interpreted as a pathway activation score.

### Data and code availability

The raw FASTQ files whole-exome sequencing data generated in this study will be submitted to the European Genome-phenome Archive (EGA). Processed single-cell RNA sequencing data generated in this study will be uploaded to the Gene Expression Omnibus (GEO) and will be publicly available at the date of publication.

## Acknowledgments

We thank patients who consented to participate to the study and all clinicians, pathologists, technicians, and study nurses involved in the samples acquisition. We acknowledge the IHC facility of the Institute of Pathology (Petra Hirschmann), as well as the microscopy (Ewelina Bartoszek, Loic Sauteur) and bioinformatic facilities (Robert Ivanec, Julien Roux, Florian Geier) of the DBM for technical and scientific support. We thank the Next Generation Sequencing Platform of the University of Bern for performing the scRNA sequencing. We are grateful to Yu Chen (MSKCC, US) for kindly providing the MSK-PCa2 organoid line, to Christopher McGinnis and Zev Gartner (UCSF) for providing MULTI-seq reagents, to Ilaria Alborelli and Ester Cannizzaro for performing the targeting sequencing, and to Salvatore Piscuoglio for support with WES analysis. Financial support was provided by funding from Krebsliga beider Basel (KLbB-5329-03-2021 to CL), the Swiss National Science Foundation (320030_205086 to CL), and the Department of Surgery of the University Hospital Basel (*PMC* Platform to CL). Components of several figures were created with BioRender.com.

## Author contributions

C.L., R.P., R.D., and R.S. conceived the study. C.L. supervised the study. R.P., R.D., R.S., J.W., R.M., D.B., and Z.D. designed and conducted experiments. R.P., R.D., R.S., and L.R. analyzed data and performed data visualization. H.P., T.V., F.S., A.J.T., H.S., C.A.R., L.B. contributed to patient enrollment, collection of samples and clinical data, and provided clinical advices. T.V. and L.B. performed the histopathological evaluation. N.A. and J-P.T supervised animal experiments and provided experimental support. C.L., R.P., and R.D. wrote the initial manuscript and all the authors contributed to the final version.

