## Supplementary Material for "A modular patient-derived organoid–xenograft platform reveals molecular and clinical trajectories of prostate cancer progression"

#### Supplementary Material List

##### Supplementary Figures

Figure S1. Phenotypic and molecular characterization of P20-11 and P20-23 model series

Figure S2. Single-cell transcriptomic profiles of P20-11 and P20-23 model series

Figure S3. Phenotypic characterization of patient derived organoid xenograft (PDOX) models

Figure S4. Dose-response analysis

Figure S5. Lack of *AR* mutation in diagnostic biopsy of patient P20-23 and glucocorticoid treatment replicate

Figure S6. Single-cell RNA-sequencing analysis of P20-11 patient-derived organoids under androgen deprivation

Figure S7. Phenotypic characterization of P20-11 organoids and matched patients' samples

Figure S8. Expression patterns of PROX1 and ALDH1A1 in NEPC and DNPC patient samples

##### Supplementary Tables

Table S1. List of PDO, PDOX, and PDOXO models used for analyses

Table S2. Quantification of AR, PROX1 and ALDH1A1 positive cells in PDOXOs

Table S3. List of antibodies used in the study

##### Supplementary Dataset

Supplementary dataset 1

### Figure S1

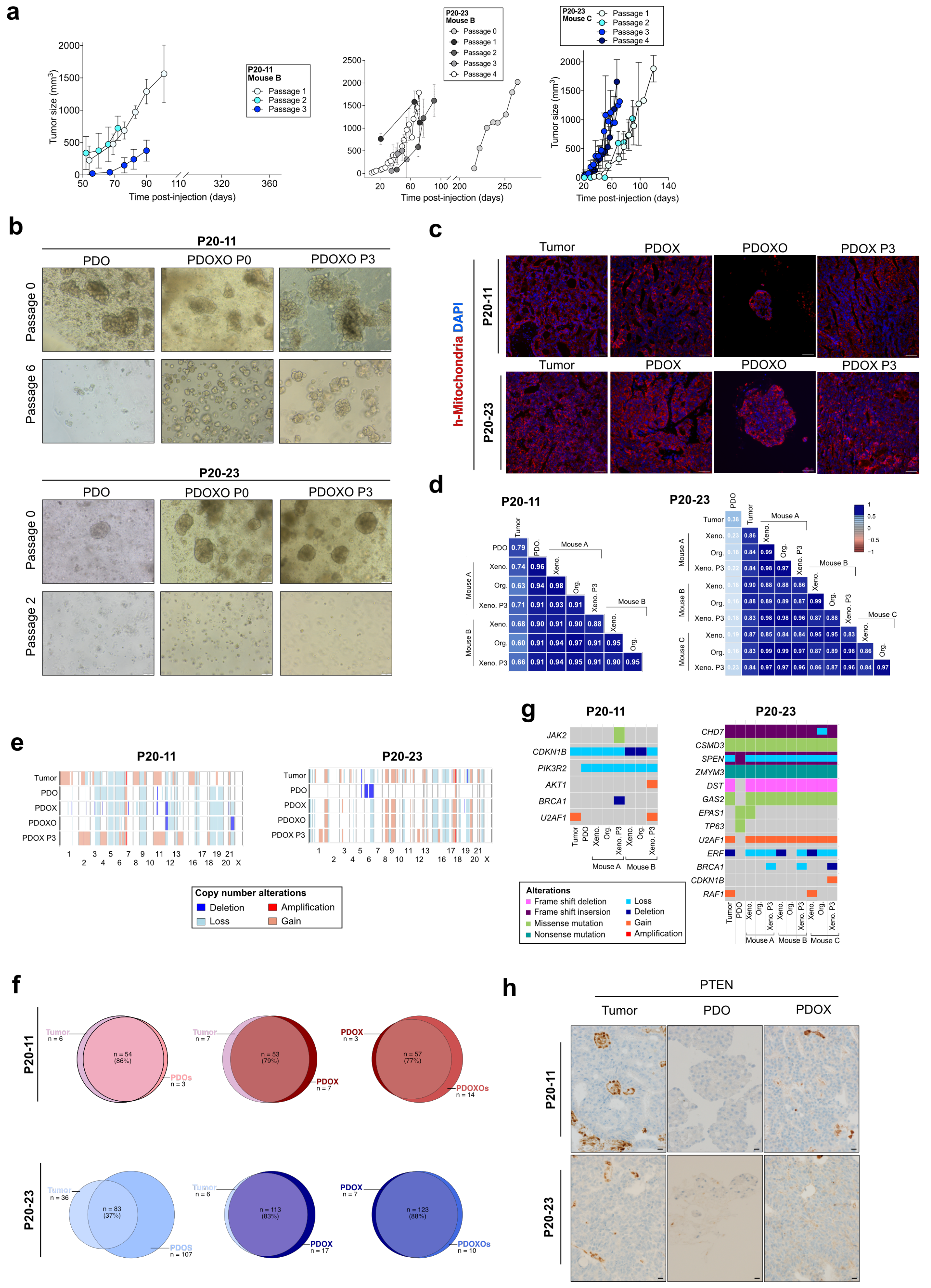

a
P20-11 Mouse B
Passage 1
Passage 2
Passage 3
Tumor size (mm3)
Time post-injection (days)
P20-23 Mouse B
Passage 0
Passage 1
Passage 2
Passage 3
Passage 4
P20-23 Mouse C
b
P20-11
PDO
PDOXO P0
PDOXO P3
Passage 0
Passage 6
P20-23
Passage 2
c
h-Mitochondria DAPI
Tumor
PDOX
PDOXO
PDOX P3
P20-11
P20-23
d
Mouse A
Mouse B
Mouse C
Xeno.
Org.
Xeno. P3
e
Copy number alterations
Deletion
Amplification
Loss
Gain
f
Tumor
PDOs
PDOX
PDOXOs
g
JAK2
CDKN1B
PIK3R2
AKT1
BRCA1
U2AF1
CHD7
CSMD3
SPEN
ZMYM3
DST
GAS2
EPAS1
TP63
ERF
RAF1
Alterations
Frame shift deletion
Frame shift insertion
Missense mutation
Nonsense mutation
Loss
Deletion
Gain
Amplification
h
PTEN

**Figure S1. Phenotypic and molecular characterization of P20-11 and P20-23 model series. (a)** PDOX tumor growth kinetics of P20-11 and P20-23 through *in vivo* passages. Mouse sublines are indicated in the legend box of each panel. **(b)** Representative brightfield images of P20-11 and P20-23 PDOs and PDXOs at various passages *in vivo* and *in vitro*. Scale bars represent 100µm **(c)** Representative immunofluorescence staining and imaging using antibodies recognizing human mitochondria for P20-11 and P20-23. Scale bars represent 50 µm. **(d)** Pearson correlation scores between P20-11 and P20-23 model series, based on copy number variation (CNV) data inferred by Whole Exome Sequencing (WES). **(e)** CNV map of P20-11 (PDOX-B) and P20-23 (PDOX-A) model series. **(f)** Venn diagram depicting the number of shared mutations among patient tumors and derived PDOs, between PDOX tumors and PDOXOs for both P20-11 (PDOX-B) and P20-23 (PDOX-A) series. The quantification of shared mutations is based on synonymous and non-synonymous mutations detected by WES. Numbers indicate sample specific and shared mutations. Percentages in brackets represent the proportion of shared mutation between the groups. **(g)** Oncoplot depicting model-specific alterations for P20-11 and P20-23 patient tumors, PDOs, PDOXs, PDOXOs, at various *in vivo* passages. Xeno: Xenograft, Org: Organoids. (**h)** Immunohistochemistry staining with antibodies recognizing PTEN for P20-11 and P20-23 patient tumors, PDOs, and PDOXOs. Scale bars represent 20 µm.

### Figure S2

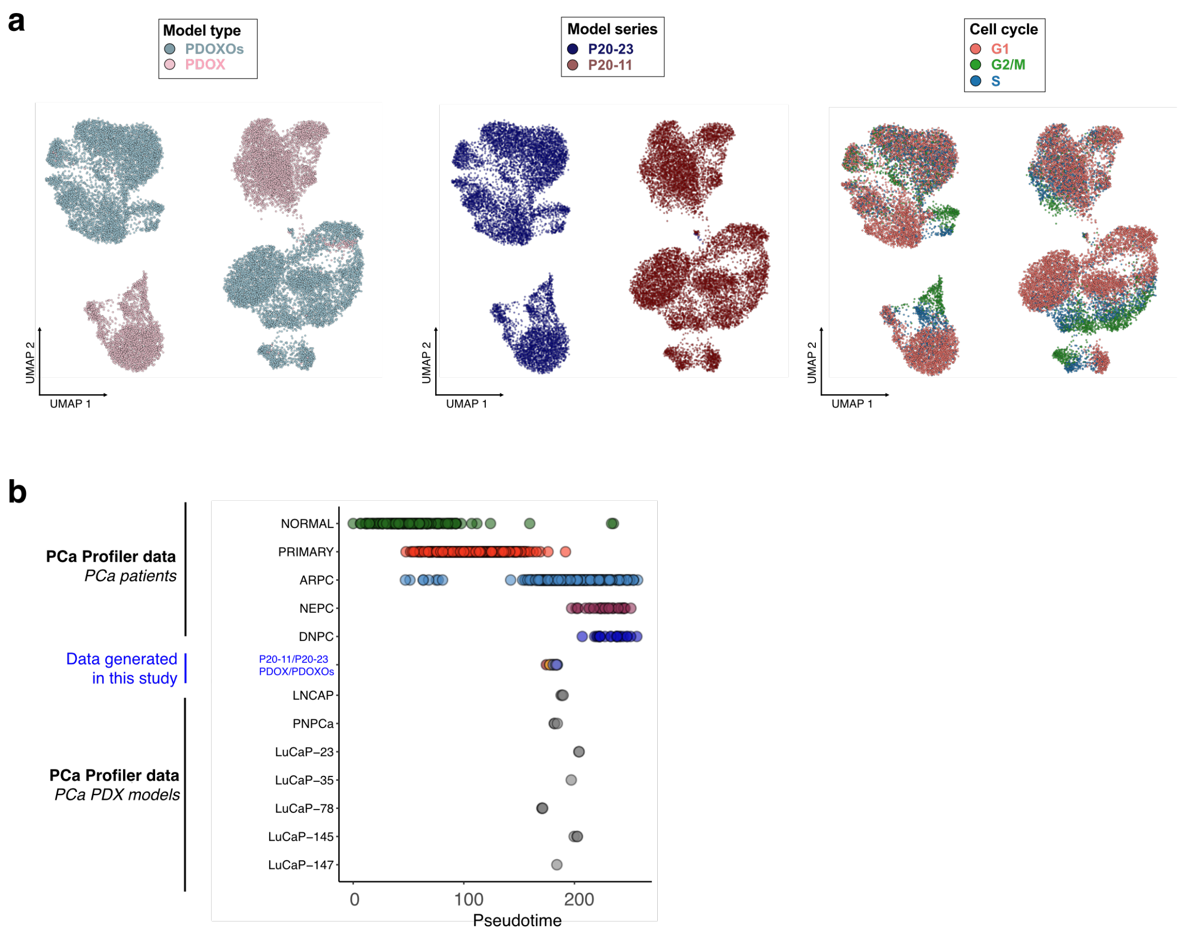


**Figure S2. Single-cell transcriptomic profiles of P20-11 and P20-23 model series.**
**(a)** UMAP projection of P20-11 and P20-23 model series color/coded by associated metadata. From left to right: model type, model series, cell cycle phase. **(b)** Dotplot representing pseudotime values of PCa patient samples and PCa experimental models from the Prostate Cancer Atlas. Each dot represents an individual sample ranked according to its pseudotime value, as calculated using the PCa Atlas web interface. Single-cell data generated in this study were aggregated by summing read counts and positioned within the different molecular subtypes and model categories available in the Prostate Cancer Atlas tool.

### Figure S3

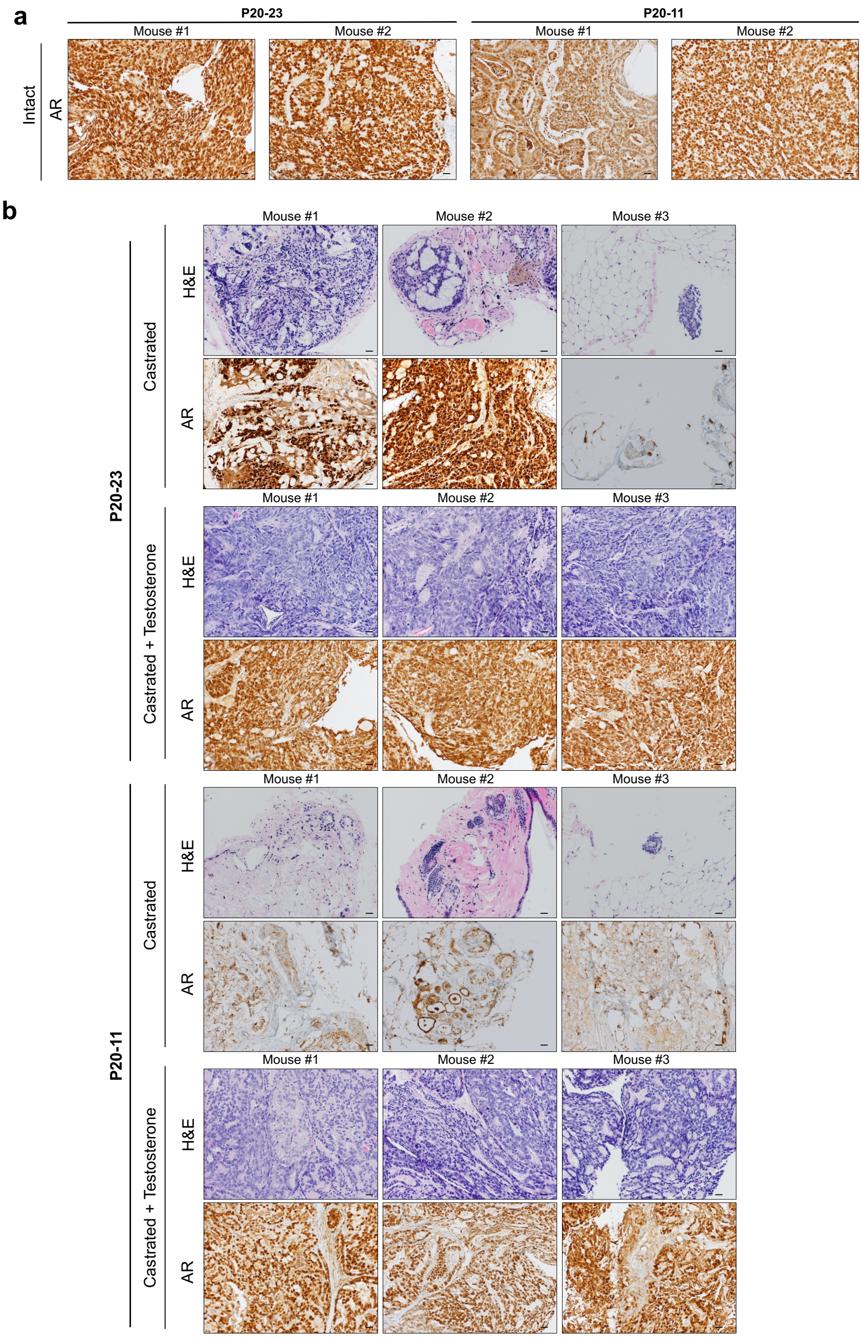


**Figure S3. Phenotypic characterization of patient-derived organoid xenograft (PDOX) models**. **(a)** Immunohistochemistry staining with antibodies recognizing the androgen receptor (AR) in P20-11 and P20-23 intact PDOXs (*i.e.* not subjected to castration). Scale bars represent 20 µm. **(b)** H&E and Immunohistochemistry analysis with antibodies recognizing the androgen receptor (AR) in P20-23 and P20-11 PDOXs following castration and post-testosterone rescue (*i.e*. testosterone implantation post-castration). Scale bars represent 20 µm.

### Figure S4

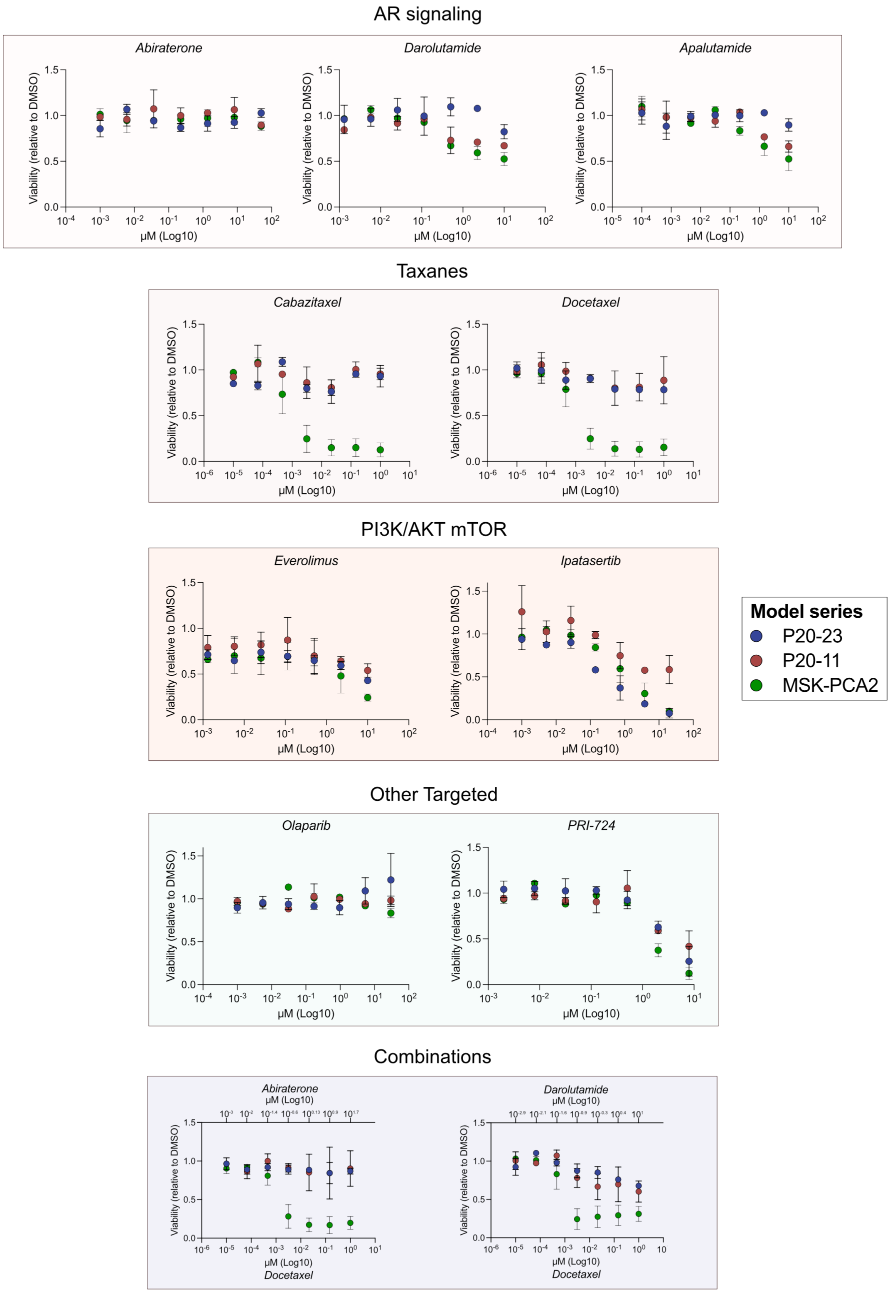


**Figure S4. Dose-response analysis.** Drug dose response analysis for the indicated drugs and models (*i.e.* P20-11 and P20-23 PDOXOs series, PDOs for MSK-PCa2). Data are represented as means relative to the vehicle control and error bars show standard deviation. Two independent biological replicates were performed. For drug combination experiments, two x-axes are shown to indicate the concentrations applied for each compound.

#### Figure S5

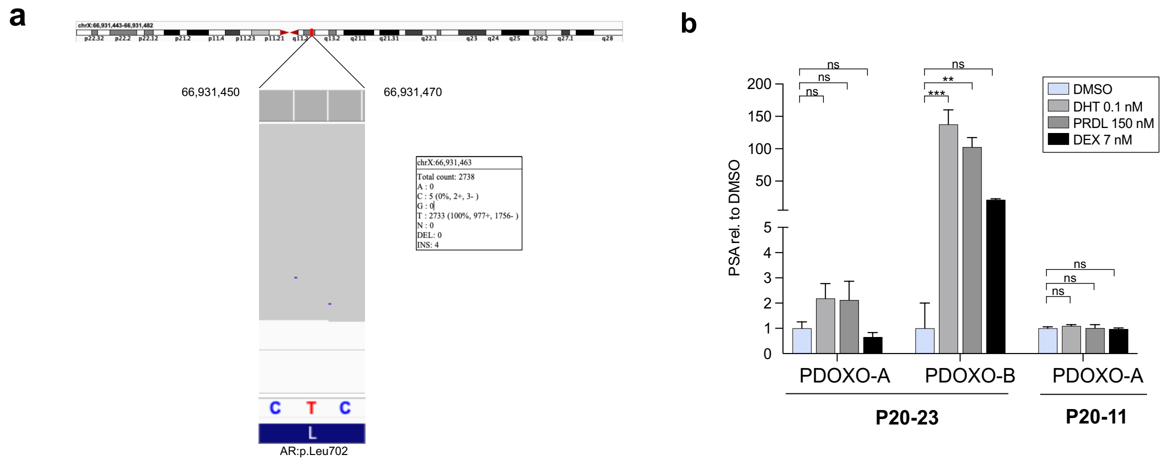


**Figure S5. Lack of AR mutation in diagnostic biopsy of patient P20-23 and glucocorticoid treatment replicate**. **(a)** Genomic sequencing using the HRR-Prostate targeted panel in the initial diagnostic biopsy. Inset highlights the codon in exon 4 of the Androgen Receptor (AR) locus encoding for L702, showing the absence of mutation at this site. **(b)** PSA levels normalized to total protein content shown as relative to DMSO (DHT 0 + vehicle) upon DHT 0.1 nM, prednisolone (150 nM) or dexamethasone (7 nM) 5-day exposure and measured by ELISA in the culture medium. Statistical test for P20-23 (PDOX-A): One-way Anova, multiple comparisons to DHT 0 nM, adjusted P values: DHT 0.1 nM = 0.1031, PRDL = 0.2369, DEX = 0.6148. Statistical test for P20-23 (PDOX-B): One-way Anova, multiple comparisons to DHT (0 nM), adjusted P values: DHT (0.1 nM) = 0.0002, PRDL = 0.0018, DEX = 0.6068. Statistical test for P20-11: One-way Anova, multiple comparisons to DHT (0 nM), adjusted P values: DHT (0.1 nM) = 0.6418, PRDL > 0.9999, DEX = 0.9590. Statistical tests were performed on values normalized to total protein content and non-normalized to DMSO. Replicate 2 is represented and replicate 1 can be found in Figure 3.

### Figure S6

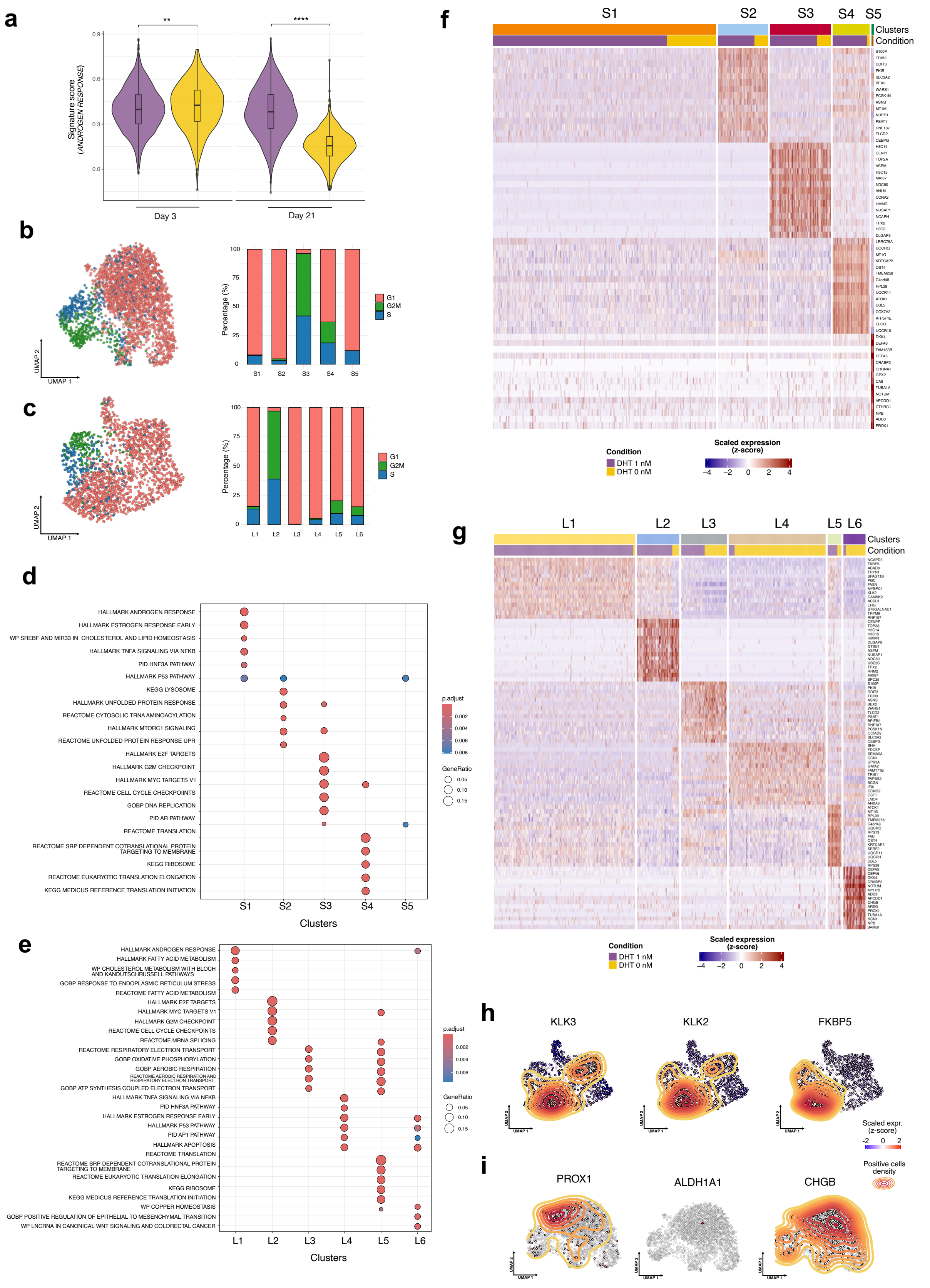
a
Signature score (ANDROGEN RESPONSE)
**
****
Day 3
Day 21
b
UMAP 2
UMAP 1
Percentage (%)
G1
G2M
S
S1 S2 S3 S4 S5
c
L1 L2 L3 L4 L5 L6
d
HALLMARK ANDROGEN RESPONSE
HALLMARK ESTROGEN RESPONSE EARLY
WP SREBF AND MIR33 IN CHOLESTEROL AND LIPID HOMEOSTASIS
HALLMARK TNFA SIGNALING VIA NFKB
PID HNF3A PATHWAY
HALLMARK P53 PATHWAY
KEGG LYSOSOME
HALLMARK UNFOLDED PROTEIN RESPONSE
REACTOME CYTOSOLIC TRNA AMINOACYLATION
HALLMARK MTORC1 SIGNALING
REACTOME UNFOLDED PROTEIN RESPONSE UPR
HALLMARK E2F TARGETS
HALLMARK G2M CHECKPOINT
HALLMARK MYC TARGETS V1
REACTOME CELL CYCLE CHECKPOINTS
GOBP DNA REPLICATION
PID AR PATHWAY
REACTOME TRANSLATION
REACTOME SRP DEPENDENT COTRANSLATIONAL PROTEIN TARGETING TO MEMBRANE
KEGG RIBOSOME
REACTOME EUKARYOTIC TRANSLATION ELONGATION
KEGG MEDICUS REFERENCE TRANSLATION INITIATION
Clusters
p.adjust
GeneRatio
e
HALLMARK ANDROGEN RESPONSE
HALLMARK FATTY ACID METABOLISM
WP CHOLESTEROL METABOLISM WITH BLOCH AND KANDUTSCHRUSSELL PATHWAYS
GOBP RESPONSE TO ENDOPLASMIC RETICULUM STRESS
REACTOME FATTY ACID METABOLISM
HALLMARK E2F TARGETS
HALLMARK MYC TARGETS V1
HALLMARK G2M CHECKPOINT
REACTOME CELL CYCLE CHECKPOINTS
REACTOME MRNA SPLICING
REACTOME RESPIRATORY ELECTRON TRANSPORT
GOBP OXIDATIVE PHOSPHORYLATION
GOBP AEROBIC RESPIRATION
REACTOME AEROBIC RESPIRATION AND RESPIRATORY ELECTRON TRANSPORT
GOBP ATP SYNTHESIS COUPLED ELECTRON TRANSPORT
HALLMARK TNFA SIGNALING VIA NFKB
PID HNF3A PATHWAY
HALLMARK ESTROGEN RESPONSE EARLY
HALLMARK P53 PATHWAY
PID AP1 PATHWAY
HALLMARK APOPTOSIS
REACTOME TRANSLATION
REACTOME SRP DEPENDENT COTRANSLATIONAL PROTEIN TARGETING TO MEMBRANE
REACTOME EUKARYOTIC TRANSLATION ELONGATION
KEGG RIBOSOME
KEGG MEDICUS REFERENCE TRANSLATION INITIATION
WP COPPER HOMEOSTASIS
GOBP POSITIVE REGULATION OF EPITHELIAL TO MESENCHYMAL TRANSITION
WP LNCRNA IN CANONICAL WNT SIGNALING AND COLORECTAL CANCER
Clusters
f
Clusters
Condition
DHT 1 nM
DHT 0 nM
Scaled expression (z-score)
g
h
KLK3
KLK2
FKBP5
Scaled expr. (z-score)
i
PROX1
ALDH1A1
CHGB
Positive cells density

**Figure S6. Single-cell RNA-sequencing analysis of P20-11 patient-derived organoids under androgen deprivation. (a)** Violin and boxplots illustrating the signature score for *ANDROGEN RESPONSE* (MsigDB, Hallmark) at day 3 and day 21, colored by culture conditions. wo-tailed t-tests (after verification of normality) were used for pairwise comparisons (*, **, *** and **** indicate adjusted P values < 0.01, 0.001, 0.0001, and 0.00001, respectively). Adjusted P values: day 3 = 0.00247, day 21 = 1.67e-238. **(b-c)** UMAP projections showing cell cycle phase distribution and bar plots indicating the percentage of cells in each phase per cluster at day 3 **(b)** and day 21 **(c)**. **(d-e)** Dot plot showing Top 5 GSEA-enriched pathways (MsigDB, Hallmark & GOBP collections) in each cluster at day 3 **(d)** and Day 21 **(e)**. Dot color: adjusted P value, dot size: Gene Ratio (fraction of genes in the pathway). **(f-g)** Heatmap illustrating the scaled (z-score) expression profiles of the top 15 cluster-specific marker genes, identified by FindAllMarkers (min.pct = 0.5, log2FC = 0.5) at day 3 **(f)** or day 21 **(g)**. Culture conditions are shown in the top annotation tracks. **(h-i)** UMAP feature plot with density overlay on cells at day 3 **(h)** and day 21 **(i)**. Cells are color-coded according to their scaled expression for indicated markers, and density overlay of cells with a positive scaled expressed is shown as red line

### Figure S7

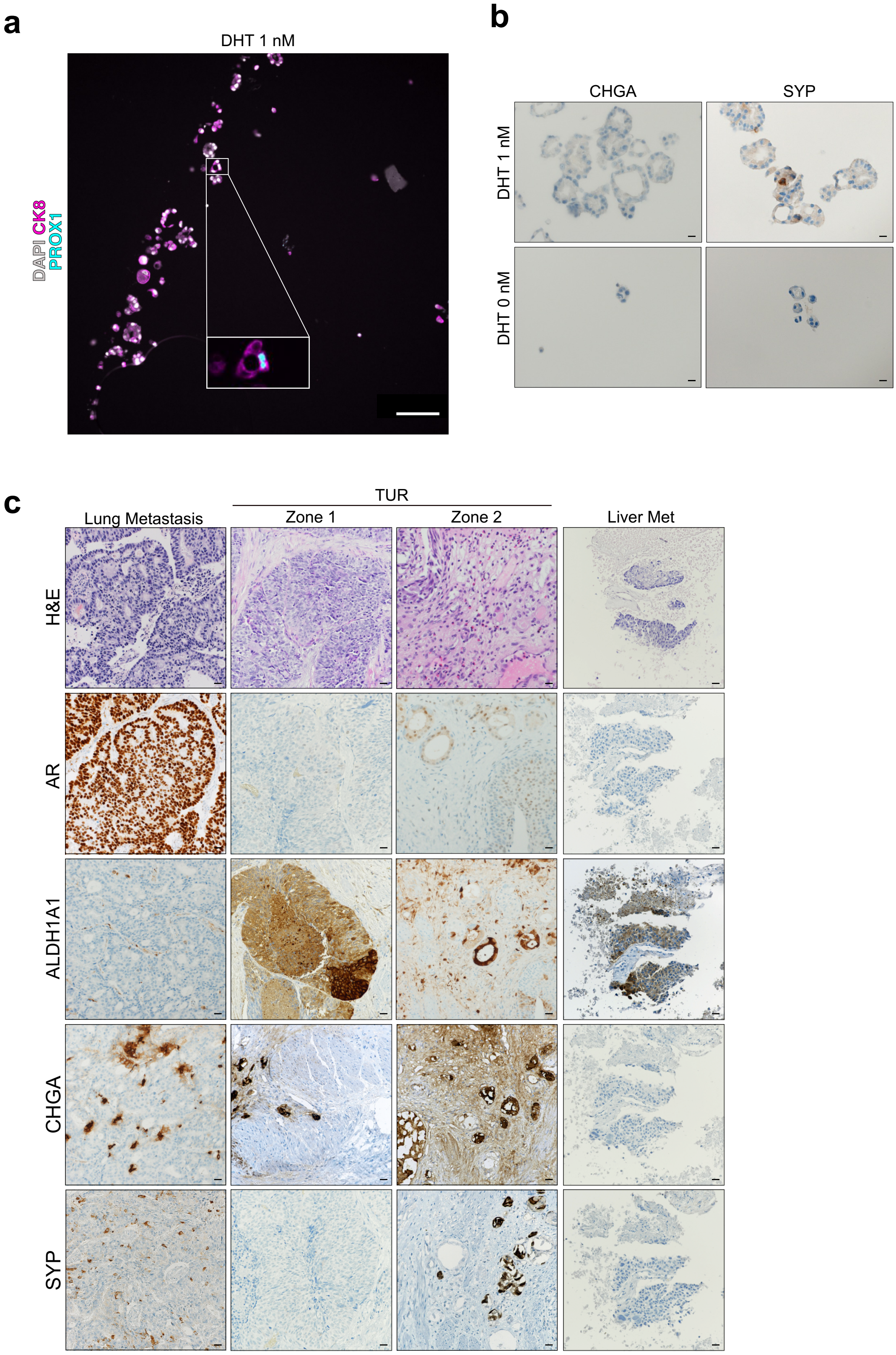
a
DHT 1 nM
DAPI CK8 PROX1
b
CHGA
SYP
DHT 1 nM
DHT 0 nM
c
TUR
Lung Metastasis
Zone 1
Zone 2
Liver Met
H&E
AR
ALDH1A1
CHGA
SYP

**Figure S7. Phenotypic characterization of P20-11 organoids and matched patients' samples. (a)** Immunofluorescence of P20-11 PDOXOs cultured in DHT 1 nM (CTRL) for 42 days. PDOXOs were stained with antibodies recognizing cytokeratin 8 (CK8) and PROX1. The image highlights the scarcity of PROX1+ cells in the DHT proficient condition (1 nM) and is a higher magnification of the image shown in Figure 4e. The scale bar represents 50 µm. **(b)** Immunohistochemistry staining with antibodies recognizing Chromogranin A (CHGA) and Synaptophysin (SYP) of P20-11 PDOXOs cultured in either DHT 1 nM (CTRL) and DHT 0 nM (ADT) conditions for 42 days. Scale bars represent 50µm. **(c)** Phenotypic characterization of initial P20-11 lung metastasis sample and progression samples (*i.e.* TUR and liver metastasis). Top: Hematoxylin and eosin staining. Bottom: Immunohistochemistry staining with antibodies recognizing the androgen receptor (AR), ALDH1A1, chromogranin A (CHGA) and synaptophysin (SYP). Some images are already represented in Figure 4g but are also included here to visualize the full phenotypic profile of all samples. Scale bars represent 20 µm.

### Figure S8

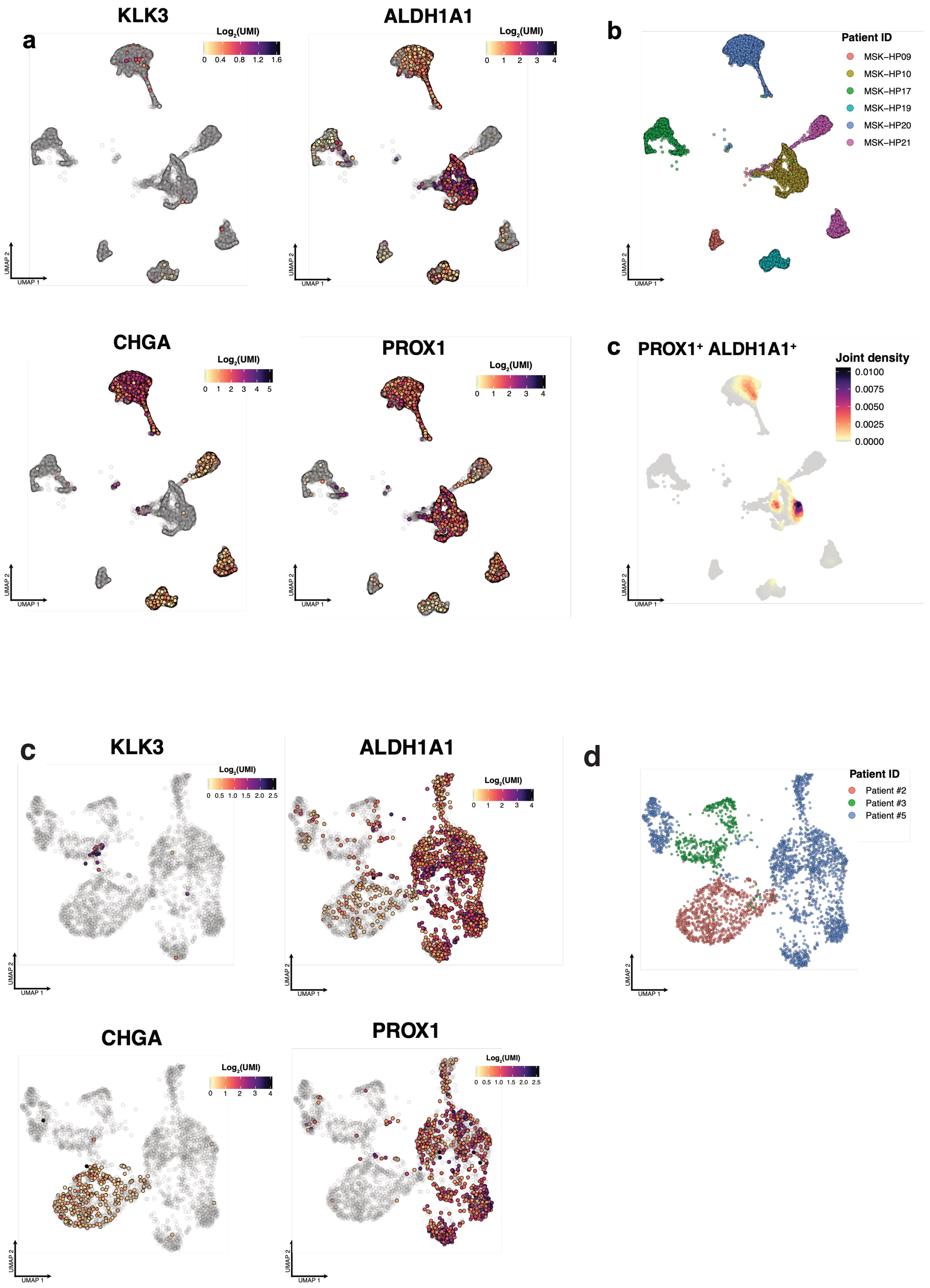
a
KLK3
ALDH1A1
CHGA
PROX1
Log2(UMI)
UMAP 1
UMAP 2
b
Patient ID
MSK–HP09
MSK–HP10
MSK–HP17
MSK–HP19
MSK–HP20
MSK–HP21
c
PROX1+ ALDH1A1+
Joint density
0.0100
0.0075
0.0050
0.0025
0.0000
c
KLK3
ALDH1A1
CHGA
PROX1
d
Patient ID
Patient #2
Patient #3
Patient #5

**Figure S8. Expression patterns of PROX1 and ALDH1A1 in NEPC and DNPC patient samples. (a-b)** UMAP projection of single-cell RNA-seq data from neuroendocrine prostate cancer (NEPC) and double-negative prostate cancer (DNPC) samples from the Zaidi et al. dataset [42]. Cells are color-coded by expression ($log_2$(UMI)) of KLK3, CHGA, ALDH1A1, and PROX1 **(a)** and sample distribution **(b)**. **(c-d)** UMAP projection of single-cell RNA-seq data from NEPC and DNPC samples from the Dong et al. dataset [41]. Cells are color-coded by expression ($log_2$(UMI)) of KLK3, CHGA, ALDH1A1, and PROX1 **(c)** and sample distribution **(d)**. Only patients with a $CHGA^+$ phenotype (NEPC) or a $KLK3^-$/$CHGA^-$ phenotype (DNPC) were included in the integrated analysis.

#### Table S2. Quantification of AR, PROX1 and ALDH1A1 positive cells in PDOXOs

| PROX1 | | |
|---|---|---|
| **Condition (DHT, nM)** | **Positive** | **Negative** |
| 0 | 2270 | 6440 |
| 1 | 80 | 7370 |
| **ALDH1A1** | | |
| **Condition (DHT, nM)** | **Positive** | **Negative** |
| 0 | 61 | 43 |
| 1 | 24 | 1722 |
| **AR** | | |
| **Condition (DHT, nM)** | **Positive** | **Negative** |
| 0 | 36 | 26 |
| 1 | 1976 | 50 |

**Table S3. List of antibodies used in the study**

| Primary antibodies used for IHC | | | | |
|---|---|---|---|---|
| **Marker** | | **Reference** | **Manufacturer** | **Dilution** |
| ALDH1A1 | | ab52492 | Abcam | 1:50 |
| Androgen Receptor | | 760-4605 | Ventana | Ready-to-use |
| Chromogranin A | | 760-2519 | Ventana | Ready-to-use |
| ERG | | 790-4576 | Ventana | Ready-to-use |
| PSMA | | 760-6076 | Ventana | Ready-to-use |
| PTEN | | 790-5097 | Ventana | Ready-to-use |
| Synaptophysin | | | | |
| **Primary antibodies used for IF** | | | | |
| **Marker** | | **Reference** | **Manufacturer** | **Dilution** |
| Androgen Receptor | | ab133273 | Abcam | 1:200 |
| Cytokeratin 5 | | 905901 | Biolegend | 1:1000 |
| Cytokeratin 8 | | 904801 | Biolegend | 1:1000 |
| Human Mitochondria | | ab92824 | Abcam | 1:2000 |
| PROX1 | | 14963T | BioConcept | 1:500 |
| **Secondary Antibodies used for IF** | | | | |
| **Antigen** | **Fluorochrome** | **Reference** | **Manufacturer** | **Dilution** |
| Goat anti-chicken IgY | Alexa Fluor® 488 | A-21437 | Invitrogen | 1:1000 |
| Goat anti-rabbit IgG | Alexa Fluor® 555 | A-21428 | Invitrogen | 1:1000 |
| Goat anti-mouse IgG | Alexa Fluor® 647 | A-21237 | Invitrogen | 1:1000 |
